# Human organoid systems reveal *in vitro* correlates of fitness for SARS-CoV-2 B.1.1.7

**DOI:** 10.1101/2021.05.03.441080

**Authors:** Mart M. Lamers, Tim I. Breugem, Anna Z. Mykytyn, Yiquan Wang, Nathalie Groen, Kèvin Knoops, Debby Schipper, Jelte van der Vaart, Charlotte D. Koopman, Jingshu Zhang, Douglas C. Wu, Petra B. van den Doel, Theo Bestebroer, Corine H. GeurtsvanKessel, Peter J. Peters, Mauro J. Muraro, Hans Clevers, Nicholas C. Wu, Bart L. Haagmans

## Abstract

A new phase of the COVID-19 pandemic has started as several SARS-CoV-2 variants are rapidly emerging globally, raising concerns for increased transmissibility. As animal models and traditional *in vitro* systems may fail to model key aspects of the SARS-CoV-2 replication cycle, representative *in vitro* systems to assess variants phenotypically are urgently needed. We found that the British variant (clade B.1.1.7), compared to an ancestral SARS-CoV-2 clade B virus, produced higher levels of infectious virus late in infection and had a higher replicative fitness in human airway, alveolar and intestinal organoid models. Our findings unveil human organoids as powerful tools to phenotype viral variants and suggest extended shedding as a correlate of fitness for SARS-CoV-2.

**One-Sentence Summary:** British SARS-CoV-2 variant (clade B.1.1.7) infects organoids for extended time and has a higher fitness *in vitro*.

## Introduction

The ongoing COVID-19 pandemic is causing an immense global health crisis, affecting all parts of society. As many countries have started the rollout of vaccines, several SARS-CoV-2 variants have emerged that raise concerns for increased transmissibility and escape from immunity (*1*). The B.1.1.7 variant, also known as the British variant or VOC-202012/01, now has become the dominant variant in Europe, and is increasing in frequency in Asia and the American continent (*2*). Epidemiological data suggest that this variant is 35-100% more transmissible than the ancestral lineage and associated with higher viral loads (*3–6*). As epidemiological studies are easily confounded by public health measures or founder effects, they should preferably be complemented by experimental studies on viral fitness and infectiousness. However, correlates of fitness or infectiousness for respiratory viruses - let alone SARS-CoV-2 - are ill-defined and complicated by the lack of relevant experimental systems to compare virus strains.

Traditionally, virus strains are compared using animal models, after obtaining a clinical isolate. However, viruses with mutations in the spike protein, for example to optimize ACE2 binding, may be selected in animals (e.g. hamsters and ferrets) (*7*). In fact, mutations already present in the South African (B.1.351) and Brazilian (P.1) variants have been shown to allow the virus to infect common laboratory mice (*8*). Adaptation of the virus when grown *in vitro* poses additional challenges. SARS-CoV-2 propagation on the African green monkey cell line Vero (or derivatives, such as VeroE6) rapidly leads to culture adaptation in the spike multibasic cleavage site (MBCS), generating viruses that cannot effectively infect human airway cells or hamsters (*9–13*) thus precluding the comparison of Vero-grown SARS-CoV-2 isolates. Reverse genetic approaches could be used to slow down cell culture adaptation, but these are mainly used for comparing point mutations as cloning entirely new viruses *de novo* takes considerable time. Thus, relevant systems to rapidly compare different clinical isolates need to be explored.

Standardized SARS-CoV-2 human organoid systems may be highly relevant models for assessing differences between viral variants. Organoids are 3D *in vitro* cultures grown from stem cells which consist of organ-specific cell types that self-organize (*14, 15*). Recently, organoids have emerged as models for infectious diseases (*16–20*), including for study of SARS-CoV-2 tropism and host responses in the lung and intestine (*10, 11, 21-27*). The recent development of alveolar type 2 (AT2) cell organoids (*24–27*) enables modelling of the alveolar epithelium to SARS-CoV-2 infection. Most importantly, air-liquid interface-differentiated human airway organoid cultures may be used to propagate SARS-CoV-2 without causing cell culture adaptations (*10*). In this study, we have established a platform to compare SARS-CoV-2 isolates using human airway, alveolar and intestinal organoids. We used these systems to assess virus shedding and replicative fitness of the B.1.1.7 SARS-CoV-2 variant. Such studies are urgently needed to complement epidemiological findings and to identify correlates of fitness or infectiousness that can be measured in *in vitro* systems.

### Propagating SARS-CoV-2 B.1.1.7 without cell culture adaptations

A SARS-CoV-2 B.1.1.7 variant was isolated from an oropharyngeal sample. To prevent culture adaptation to VeroE6 cells (*10*), the virus was grown on Calu-3 cells and propagated to passage three (P3) for further experiments along with an ancestral (clade B) isolate (Bavpat-1). Deep-sequencing indicated that the obtained B.1.1.7 isolate contained all mutations associated with this clade (Table S1). Next, we confirmed that B.1.1.7 P3 did not contain culture-associated mutations in the MBCS (Fig. S1A-D) or in the rest of the genome (Fig. S1E). We also confirmed that the Bavpat-1 isolate sequence was not culture adapted (Fig. S1F-J). The P3 stock titers were characterized by plaque assay on Calu-3 and VeroE6 cells. Both the Bavpat-1 virus and B.1.1.7 virus titers were ∼10^6^ PFU/ml on Calu-3 cells (Fig. 1A), but on VeroE6 cells, the B.1.1.7 virus titer was approximately 100-fold lower (∼10^4^ PFU/ml) compared with the Bavpat-1 virus (∼10^6^ PFU/ml) (Fig. 1B-C). Plaque sizes on both Calu-3 and VeroE6 cells were smaller for the B.1.1.7 isolate compared to Bavpat-1 (Fig. 1D). The replication kinetics of both viruses were equal on Calu-3 cells but the B.1.1.7 virus was strongly attenuated on VeroE6 cells (Fig. 1E-F). This was also evident based on cytopathic effects and numbers of infected cells at day three post-infection (Fig. 1G-H). The infectivity of both viruses was quantified using the RNA copies/PFU ratio for each cell line, which confirmed that the B.1.1.7 virus had a low relative infectivity on VeroE6 cells, but not on Calu-3 cells (Fig. 1I-J). These data show that B.1.1.7 is attenuated on VeroE6 cells. However, most labs grow their SARS-CoV-2 stocks on VeroE6 cells, which could force the virus to adapt to these cells. To investigate this, we cultured both the B.1.1.7 and Bavpat-1 non-adapted P2 viruses on VeroE6 cells for 2 passages with or without fetal bovine serum (FBS) as FBS was shown to increase the frequency of MBCS deletions (*10*). Both the B.1.1.7 and the Bavpat-1 virus grown on VeroE6 cells contained major changes in the MBCS (Fig. 2A-D). No MBCS deletions were observed in this experiment, possibly because the input viruses were grown on Calu-3 cells that would select against MBCS deletions. In the B.1.1.7 virus an additional major variant in NSP6 – M164T (Wuhan-Hu-1 numbering)– was detected after VeroE6 propagation, suggesting that adaption to these cells may also occur outside of the spike protein (Fig. 2E). Plaque size and infectious titer analysis show that these stocks were indeed VeroE6-adapted, highlighting that it is critical to grow virus stocks on cells that prevent culture adaptation (e.g. Calu-3 cells) (Fig 2. F-H).

**Figure 1.**
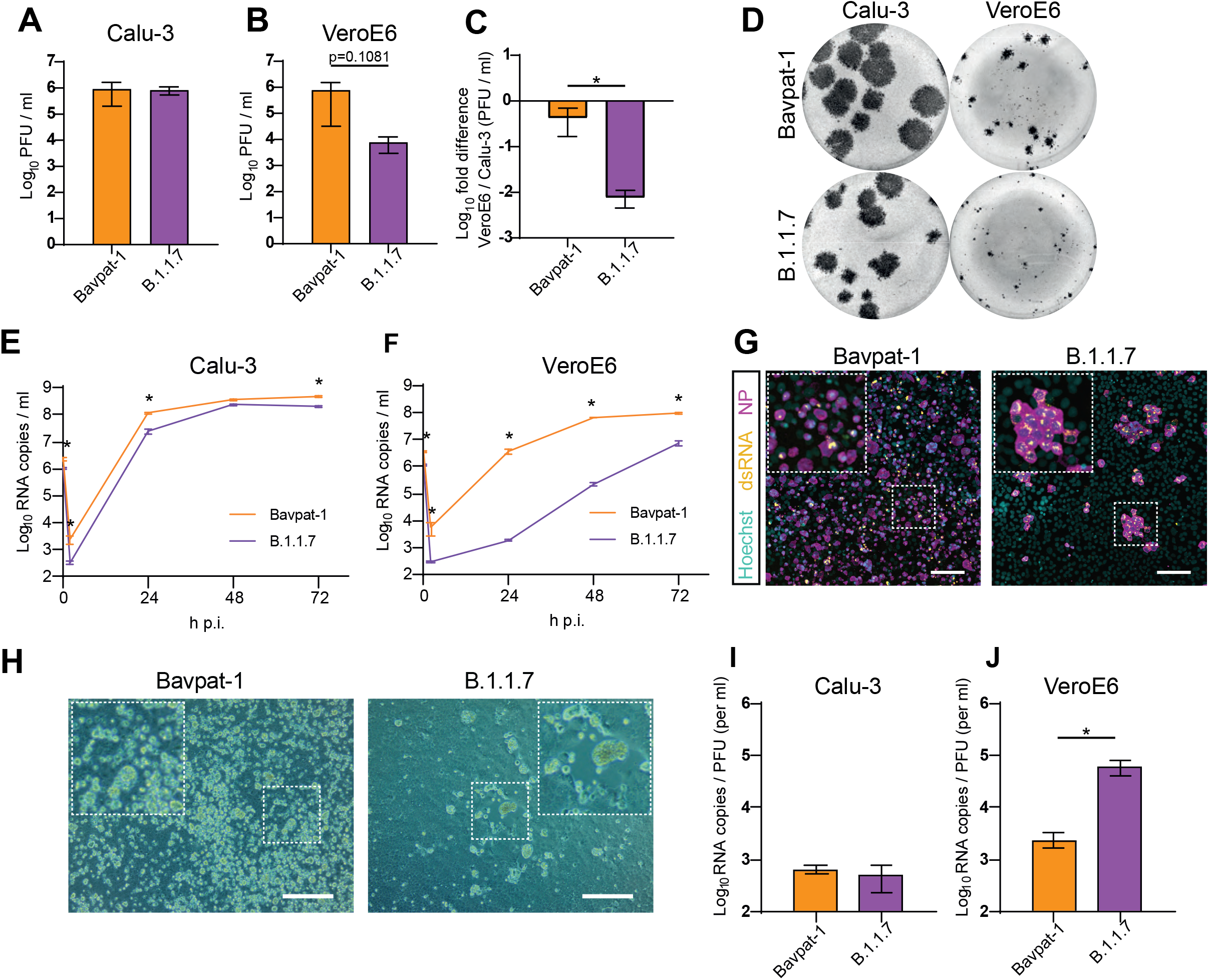
SARS-CoV-2 B.1.1.7 is attenuated on VeroE6, but not on Calu-3 cells. (A-B) Infectious virus titers (PFU/ml) of B.1.1.7 and Bavpat-1 Calu-3 P3 stocks titrated on either Calu-3 (A) or VeroE6 (B) cells. (C) Fold difference in titer (VeroE6/Calu-3) of stocks in A and B. (D) Plaque assay wells of B.1.1.7 and Bavpat-1 Calu-3 P3 viruses on Calu-3 and VeroE6 cells. (E-F) Replication kinetics of B.1.1.7 and Bavpat-1 isolates on Calu-3 (E) and VeroE6 cells (F). (G) Immunofluorescent staining of Bavpat-1 and B.1.1.7-infected VeroE6 cells at 72 hours post-infection. Viral nucleoprotein (NP, magenta), dsRNA (yellow) and hoechst (cyan) are stained. (H) Cytopathic effects induced by Bavpat-1 and B.1.1.7 in VeroE6 cells at 72 hours post-infection. (I-J) RNA/PFU ratios for Calu-3 P3 stocks of Bavpat-1 and B.1.1.7 on Calu-3 (I) and VeroE6 (J) cells. Scale bars indicate 100 µm. H p.i. = hours post-infection. Error bars indicate SEM. N=3 in all graphs. Statistical analysis was performed by unpaired T-test in A, B, C, I and J, or by two-way ANOVA in E and F. * indicates a significant difference (P<0.05).

**Figure 2.**
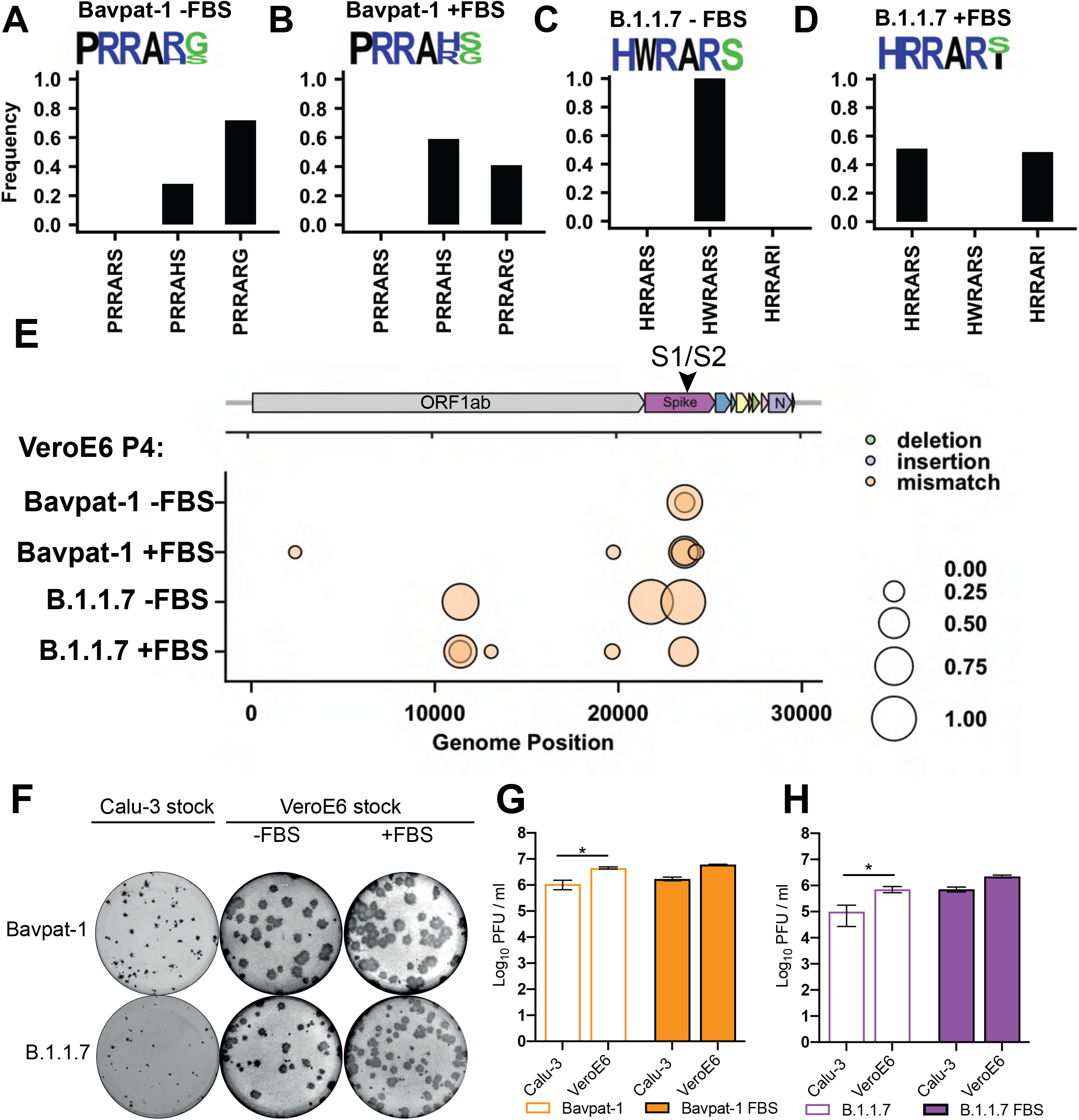
SARS-CoV-2 B.1.1.7 rapidly acquires mutations in the multibasic cleavage site and NSP6 during propagation in VeroE6 cells. (A-B) Frequency plots of multibasic cleavage site (MBCS) mutations of the Bavpat-1 VeroE6 P4 stocks without (A) and with FBS (B). (C-D) Frequency plots of MBCS mutations of the B.1.1.7 VeroE6 P4 stocks without (C) and with FBS (D). VeroE6 P4 stocks were generated by two passages on VeroE6 cells using Calu-3 P2 stocks as input viruses. MBCS plots contain the consensus MBCS amino acid sequence logo. (E) Full genome frequency plots of both Bavpat-1 and B.1.1.7 stocks. In all plots variants with a frequency >10% are depicted. (F) Plaque assay of Calu-3 P3 and VeroE6 P4 stocks on VeroE6 cells for Bavpat-1 and B.1.1.7. (G-H) Infectious virus titers (PFU/ml) of VeroE6 propagated Bavpat-1 (G) and B.1.1.7 (H) stocks with or without FBS on VeroE6 cells and Calu-3 cells. Statistical analysis was performed by unpaired T-test. * indicates a significant difference (P<0.05). Error bars indicate SEM. N=3 in G and H.

### SARS-CoV-2 B.1.1.7-infected airway organoids shed high levels of infectious virus late in infection

Humans shed infectious SARS-CoV-2 for approximately ∼10 days (*28*). However, growth kinetics cannot be studied on most cell lines for this length of time as infected cells die around 72 hours post-infection. Therefore, we assessed the replication kinetics of the B.1.1.7 and Bavpat-1 virus in air-liquid interface (ALI) human airway organoids differentiated using Pneumacult medium as described before (*22*) by sampling the organoids until day 10 post-infection from the apical side. As the RNA copies/PFU ratio of both viruses was equal on Calu-3 cells (Fig. 1I), we used viral titers determined on Calu-3 cells by plaque assay to equalize virus input at a low multiplicity of infection (moi) of 0.1. In addition, infectious virus titers in apical washes were determined on Calu-3 cells, whereas viral RNA copies were determined by qPCR. In airway organoids from different donors, the B.1.1.7 virus reached peak titers (∼10^8^ RNA copies/ml; ∼10^6^ PFU/ml) between day 2 and 4 post-infection and then continued to shed high titers (>10^7^ RNA copies/ml; >10^5^ PFU/ml) until day 10 post-infection, whereas shedding of the Bavpat-1 virus dropped by two logs between day 4 and 10 post-infection (Fig. 3A, C for RNA copies; Fig. 3B, D for infectious virus). Immunofluorescence staining on organoid wells showed that at day 3 post-infection both viruses infected roughly equal numbers of cells (Fig. 3E). However, at day 10 post-infection, cultures inoculated with B.1.1.7 still contained many infected cells, while few infected cells were observed in wells inoculated with Bavpat-1 virus (Fig. 3F). We did not observe differences in tropism: both viruses infected predominantly ciliated cells at day 3 and 10 post-infection (Fig 3G, H).

**Figure 3.**
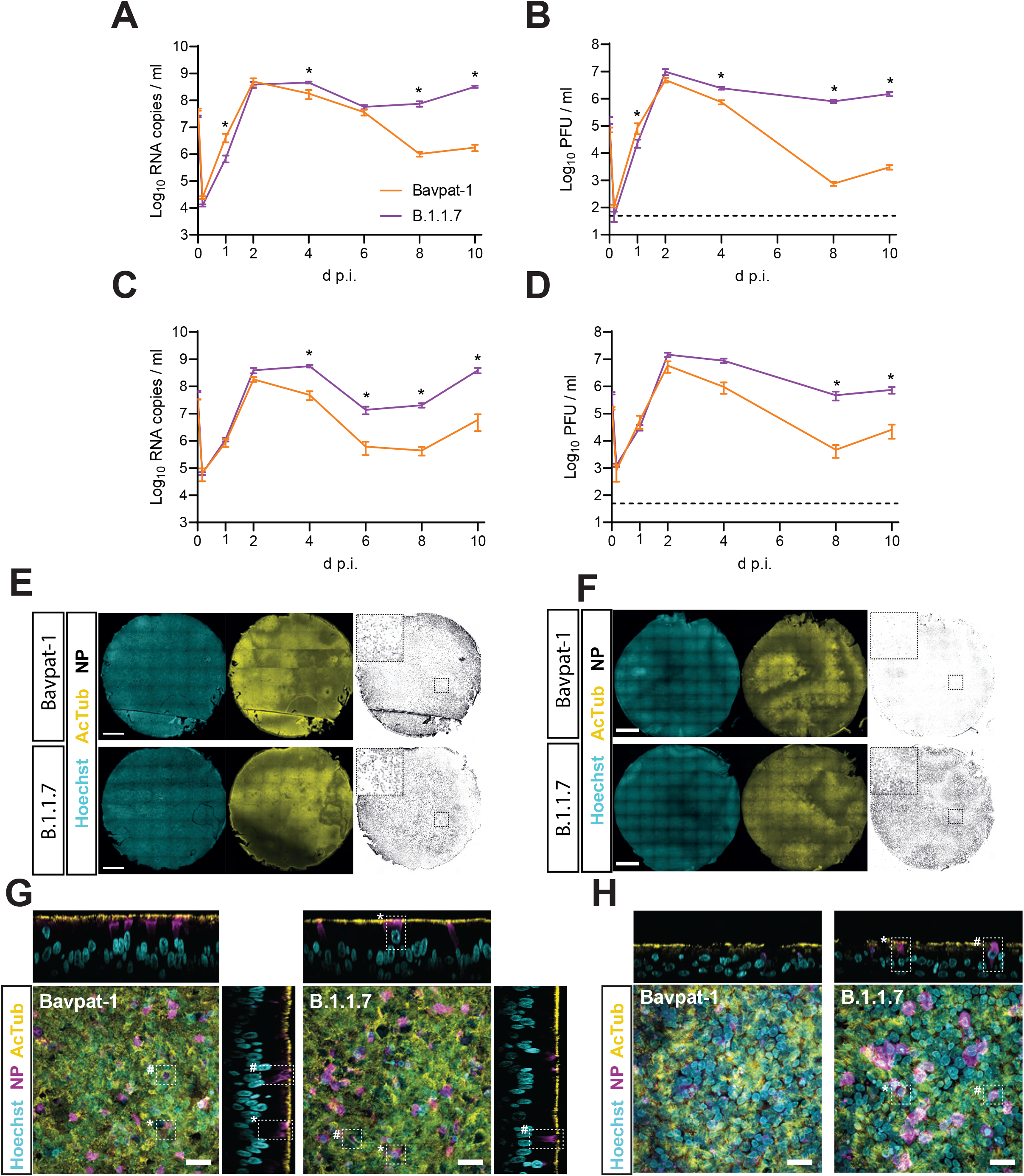
Replication kinetics and tropism of SARS-CoV-2 B.1.1.7 and Bavpat-1 in 2D differentiated human airway organoids. (A-D) Replication kinetics of Bavpat-1 and B.1.1.7 in 2D differentiated airway organoids from donor 1 (A-B) and donor 3 (C-D). Viral RNA copies (E gene) are quantified by qPCR in A and C. Infectious virus (PFU/ml) is quantified by plaque assay on Calu-3 cells in B and D. (E-F) Full well confocal images of Bavpat-1 and B.1.1.7-infected 2D airway organoids at day 3 (E) and 10 post-infection. (F). (G-H) Immunofluorescent staining of Bavpat-1 and B.1.1.7-infected 2D airway organoids at 3 d p.i. (G) and 10 d p.i. (H) showing infection of mainly ciliated cells (*) and occasionally non-ciliated cells (#). Dotted lines indicate detection limit (50 PFU/ml). Error bars indicate SEM. N=4 in A-B and N=3 in C-D. Statistical analysis was performed by two-way ANOVA. * indicates a significant difference (P<0.05).

A major limitation of the current airway organoid differentiation methods is the lack of a defined medium for differentiation. For this purpose, we established a serum-free defined airway differentiation medium based on a recipe for 3D airway differentiation (*29*). The addition of BMP2 increased ACE2 expression and ciliation, while TMPRSS2 expression and the proportion of club cells remained similar (Fig. S2A-C). Besides ciliated cells and club cells, cultures differentiated in this medium for 3-4 weeks at air-liquid interface contained goblet cells and basal cells (Fig. S2D, E). This differentiation protocol was used to investigate whether the extended shedding by B.1.1.7-infected organoids was dependent on the differentiation conditions. In these cultures - in which predominantly ciliated cells were infected (Fig. S2F) - we confirmed the extended shedding and presence of infected cells at 10 days post-infection of the B.1.1.7 virus in two donors showing that this viral phenotype is not influenced by the differentiation method (Fig. S2G, I for RNA copies; Fig. S2H, J for infectious virus; Fig. S2K for infected cells 10 dpi).

To investigate whether SARS-CoV-2 is genetically stable on 2D airway organoids, we deep-sequenced the apical washes from day 10 of the replication curves. For both the Pneumacult and BMP2 differentiated airway organoids we did not observe any indication of cell culture adaptation for Bavpat-1 and B.1.1.7 (Fig. S3; Fig. S4).

### Host responses, tropism and shedding of SARS-CoV-2 B.1.1.7-infected alveolar type 2 organoids

Whereas the human airway epithelium is highly relevant to virus transmission, alveolar cells can be used to model the severe respiratory disease caused by SARS-CoV-2. Therefore, we established human alveolar type 2 (AT2) cell organoids using a method adapted from Youk and colleagues (2020) (Fig. S5). Messenger RNA sequencing indicated that the cultured alveolar organoids expressed typical AT2 marker genes (e.g. LPCAT1, SFTPB, SFTPC), but not airway marker genes (e.g. KRT5, TP63, MUC5AC, SCGB1A1, FOXJ1) (Fig. 4A; Fig S6). Notably, AT2 cells expressed BMP4, which was shown to inhibit transdifferentiation into AT1 cells (*30*). The expression of surfactant protein C (SFTPC) was confirmed by immunofluorescent staining (Fig. 4B). AT2 organoids also contained cells that could be stained with the mature human AT2 cell surface marker antibody HTII-280 (Fig 4B). In line with the mRNA data, we did not detect any expression of the airway basal cell marker TP63 (Fig 3A; Fig. S7A-B). Transmission electron microscopy confirmed that the organoids consisted entirely of cells filled with surfactant-containing lamellar bodies (Fig. 4C; Fig. S8A-D), which formed tubular myelin structures (Fig. S8E). These cells contained apical microvilli (Fig S8F). Three dimensional AT2 organoids were susceptible to SARS-CoV-2 infection while HTII-280+ cells were the predominant cell type targeted by both B.1.1.7 and Bavpat-1 viruses (Fig. 4D, E). Next, we investigated the shedding of the B.1.1.7 and Bavpat-1 virus in these organoids after infection at a moi of 0.1. Compared to Bavpat-1, B.1.1.7 replicated to lower viral RNA titers throughout the experiment (Fig. 4F), but the infectious virus titers were slightly higher for B.1.1.7 at late time points (not significant). This discrepancy between viral RNA and infectious virus is likely to be caused by a higher peak titer for Bavpat-1 at 2 days post-infection and by the fact that in this 3D model the virus/RNA cannot be collected by daily washing of the cells. As viral spread between 3D organoids is limited, 3D systems may not be ideal for long replication kinetics experiments. Therefore, we cultured the AT2 cells in transwell inserts at 2D air-liquid interface and repeated our replication kinetics experiments (Fig. 4H-I). In this culture system, that almost exclusively contained HTII-280+ cells (Fig. S9), we observed more virus shedding of the Bavpat-1 virus in the first 3 days post-infection. However, B.1.1.7 titers remained higher than Bavpat-1 titers at later time points. Messenger RNA sequencing of the 3D AT2 organoids at day 3 post-infection showed a similar antiviral response to both Bavpat-1 and B.1.1.7 infection (Fig. 4J-K; Fig. S10).

**Figure 4.**
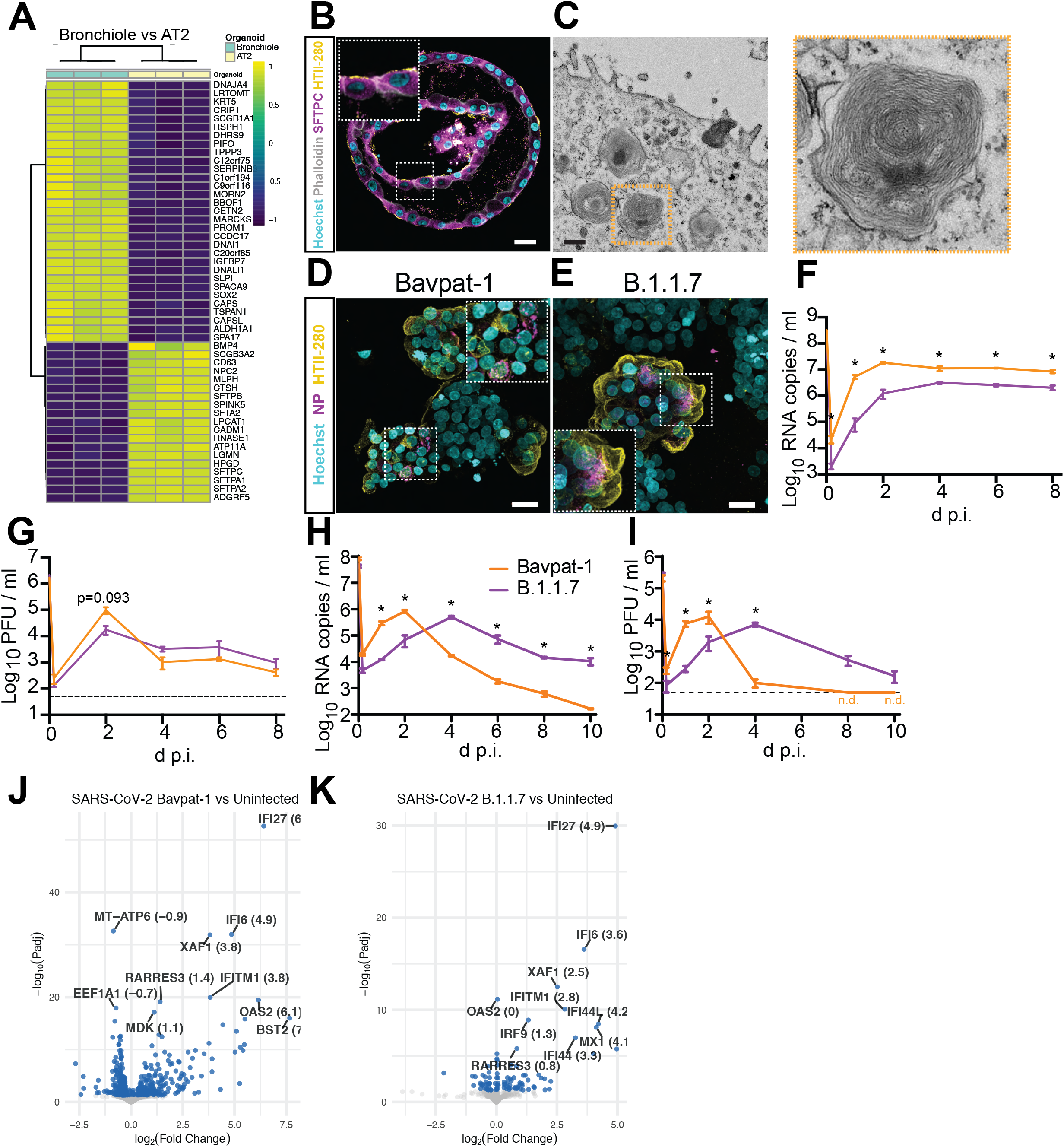
Replication kinetics, tropism and host responses of SARS-CoV-2 B.1.1.7 and Bavpat-1 in alveolar type 2 (AT2) organoids. (A) Heatmap of differentially expressed genes (DEG) comparing 2D differentiated bronchiolar and 3D AT2 organoids. Heatmap shows normalized expression of genes scaled across samples. (B) Immunofluorescent staining of an AT2 organoid with HTII-280 (yellow) and anti-surfactant protein C (SFTPC, magenta). (C) Transmission electron microscopy analysis of an AT2 cell, showing cytoplasmic lamellar bodies and apical microvilli (for a more detailed analysis see Fig. S8). (D-E) Immunofluorescent staining of AT2 organoids infected with Bavpat-1 (D) and B.1.1.7 (E) at 3 days post-infection, showing infected (nucleoprotein (NP)+; magenta) HTII-280+ AT2 cells (yellow). (F-I) Replication kinetics of Bavpat-1 and B.1.1.7 in 3D (F-G) and 2D (H-I) AT2 organoids. Viral RNA copies (E gene) are shown in F and H, while infectious virus titers (PFU/ml) are shown in G and I. (J-K) Volcano plots depicting differentially expressed genes between infected and uninfected AT2 organoids (J for Bavpat-1; K for B.1.1.7). Blue dots indicate Padj<0.05. The names of the top 10 genes with the lowest Padj are shown. In confocal images, actin is stained with phalloidin (white) and nuclei are stained with hoechst (cyan). Scale bars depict 20 µm in B and D, and 500 nm in C. Dotted lines indicate detection limit (50 PFU/ml). Error bars indicate SEM. N=3 in F-I. Statistical analysis was performed by two-way ANOVA. * indicates a significant difference (P<0.05). n.d. = not detected.

### Extended shedding of SARS-CoV-2 B.1.1.7-infected intestinal organoids

As the intestinal epithelium can also be infected by SARS-CoV-2 (*21, 31–33*), we investigated whether intestinal epithelial cells would show similar replication kinetics for B.1.1.7 compared to pulmonary cells. Similar to 3D AT2 organoids, we did not observe differences in the virus titers produced over time between both B.1.1.7 and Bavpat-1 in 3D intestinal organoids (Fig. 5A-B). In 2D intestinal cultures both viruses showed similar kinetics in the first 3 days of the infection. However, B.1.1.7 replicated to higher titers in later stages of infection compared to Bavpat-1 (Fig. 5C-D). This was evident in viral RNA levels as well as infectious virus titers. At 10 days post-infection more infected cells were observed in 2D organoids infected with B.1.1.7 compared to Bavpat-1 (Fig. 5E). Thus overall, both pulmonary and intestinal organoids cells shed high levels of infectious B.1.1.7 for a prolonged period of time.

**Figure 5.**
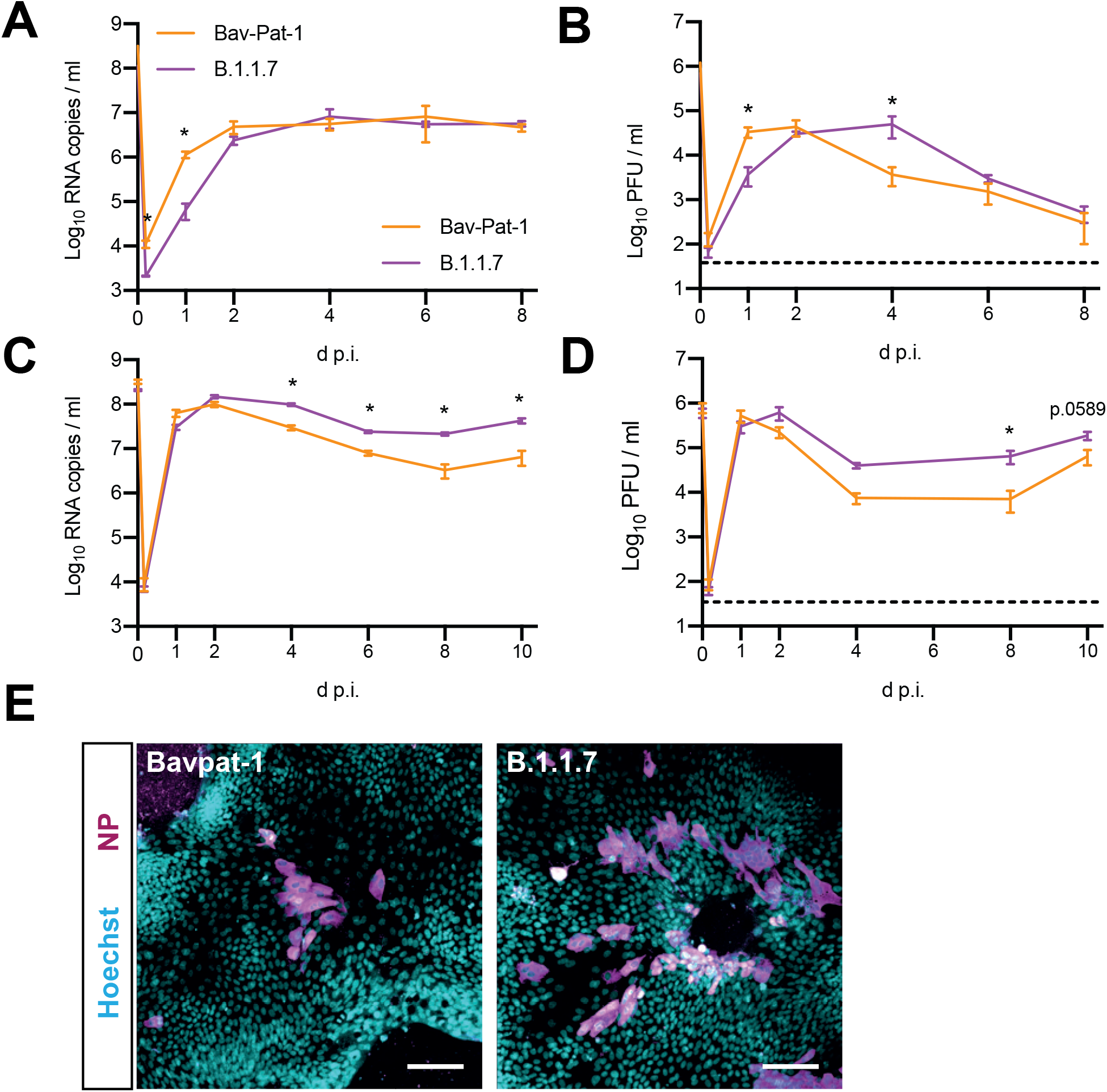
Replication kinetics of SARS-CoV-2 B.1.1.7 and Bavpat-1 in intestinal organoids. (A-D) Replication kinetics of Bavpat-1 and B.1.1.7 in 3D (A-B) and 2D (C-D) intestinal organoids. Viral RNA copies (E gene) are shown in A and C, while infectious virus titers (PFU/ml) are shown in B and D. (F) Immunofluorescent staining of viral nucleoprotein (NP; magenta) for Bavpat-1 and B.1.1.7 infected 2D intestinal organoids. Nuclei are stained with hoechst (cyan). Scale bars depict 50 µm. Dotted lines indicate detection limit (50 PFU/ml). Error bars indicate SEM. N=3 in A-B and N=4 in C-D. Statistical analysis was performed by two-way ANOVA. * indicates a significant difference (P<0.05).

### SARS-CoV-2 B.1.1.7 outcompetes Bavpat-1 in human airway organoids

As the B.1.1.7-infected cells appear to shed higher levels of virus at later stages of infection and epidemiological studies have suggested that this variant is more transmissible, we tested whether B.1.1.7 would outcompete the ancestral Bavpat-1 virus *in vitro*. For this purpose, we infected airway organoids with a mixture of Bavpat-1 and B.1.1.7 virus at a ratio of 1:1 or 5:1 (Bavpat-1:B.1.1.7; cumulative moi = 0.1). We sampled the cultures daily for 10 days and determined the relative frequency of both viruses to calculate replicative fitness as described before using Sanger sequencing and electropherogram peak analyses of a PCR fragment containing the P681H mutation (*34*). This experiment showed that B.1.1.7 outcompeted Bavpat-1 at both the 1:1 and 5:1 ratio (Fig. 6A-B for relative frequencies and C-D for replicative fitness). These findings were confirmed in independently differentiated organoids from the same donor (Fig. S11A-D), and in an independent donor (Fig. 6E). Notably, B.1.1.7 did not require more time to outcompete Bavpat-1 at the 5:1 Bavpat-1-to-B.1.1.7 ratio compared to the 1:1 ration in two separate experiments. To confirm that the fitter virus was not a recombinant we Sanger sequenced a RT-PCR fragment targeting a marker (del3675) at the other end of the genome (Fig. S11E), which confirmed the results of the P681H PCR. This was also shown by Illumina sequencing based on mapping of reads across the entire genome, which correlated well with the Sanger data (Fig. 6F). Furthermore, Sanger sequencing analysis results were confirmed using a different analysis software, the Synthego ICE indel frequency analysis tool, commonly used for analyzing CRISPR indel frequencies (Fig. S11F).

**Figure 6.**
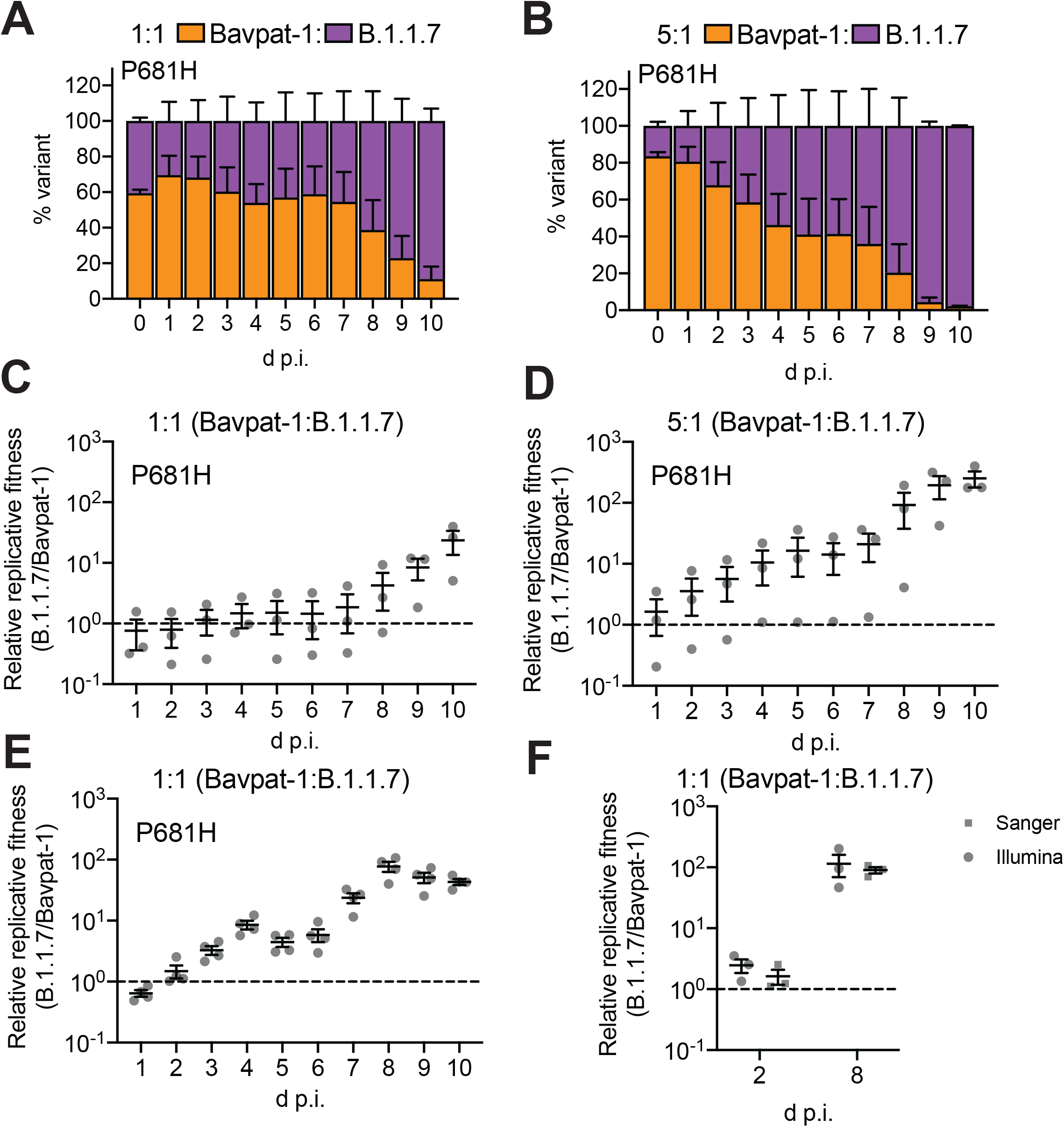
SARS-CoV-2 B.1.1.7 outcompetes Bavpat-1 in 2D differentiated human airway organoids. (A-B) Relative frequencies of Bavpat-1 and B.1.1.7 RNAs were assessed by RT– PCR and Sanger sequencing targeting the P681H mutation starting with an initial inoculum ratio of 1:1 (A) or 5:1 (B) (Bavpat-1:B.1.1.7), and a cumulative multiplicity of infection (moi) of 0.1 in airway donor 2. (C-D) These relative frequencies were used to calculate the relative replicative fitness at a 1:1 (C) and 5:1 (Bavpat-1:B.1.1.7) ratio (D). (E) Relative replicative fitness of Bavpat-1 and B.1.1.7 in airway donor 1 (starting ratio 1:1, moi 0.1). (F) Comparison of Sanger and Illumina data in estimating replicative fitness. The Sanger data is a redisplay of 3 replicates of E that were also deep-sequenced using Illumina. For the Illumina data, reads counts mapping to either B.1.1.7 or Bavpat-1 were used to calculate the replicative fitness ratio. D p.i. = days post-infection. Error bars indicate SEM.

## Discussion

Here, we established human adult-derived stem cell organoid-based systems to compare SARS-CoV-2 isolates that were propagated without cell culture adaptive mutations. We report that B.1.1.7 is strongly attenuated on VeroE6 cells, as compared with Bavpat-1. Furthermore, B.1.1.7 and Bavpat-1 rapidly acquire MBCS mutations on VeroE6 cells, but not on Calu-3 cells. Calu-3-propagated SARS-CoV-2 isolates were used for further characterization on human organoids. Our data shows that B.1.1.7-infected airway, alveolar and intestinal organoids shed high levels of infectious virus for an extended period of time. These differences were more evident in 2D systems compared with 3D systems. In addition, B.1.1.7 outcompetes Bavpat-1 in airway organoids, suggesting that B.1.1.7 has a higher fitness compared to ancestral viruses.

Clade B.1.1.7 first emerged in South England at the end of the summer in 2020. Within several months, this virus became the dominant variant in the United Kingdom and as of the 8th of March 2021 it constitutes 75.5% of cases in Europe (excluding the UK where it accounts for 98.1%), 71.7% in Africa, 47% in Oceania, 36.8% in North America (47.7% in the USA), 17.2% in Asia, 15.9% in South America, replacing locally circulating clades (*2*), suggestive of an increased transmissibility. Our results corroborate epidemiological findings that infection with the B.1.1.7 variant has been associated with a higher viral load at initial testing (*5*). In general, initial testing occurs after the 5-7 day median incubation period (*35–38*) has passed and 1-3 days after peak viral titers have been reached, indicating that our experiments accurately model the kinetics of the infection *in vivo*. A high viral load in nasal swabs from SARS-CoV-2 cases is currently the only known correlate of infectiousness for SARS-CoV-2 (*38*). *In vitro* platforms to measure correlates of infectiousness are urgently needed as countries are adapting their COVID-19 measures based on the emergence of variants that are potentially more transmissible based on scarce epidemiologic data. Another *in vitro* study that looked into the replication of B.1.1.7 and an ancestral virus on 2D primary human airway cells did not observe differences in replication, possibly because they only investigated shedding until day 3 post-infection or because their isolates were propagated on Vero cells (*39*).

It is currently unclear which mutations in the B.1.1.7 clade may cause the differences in replication kinetics and fitness observed in this study. Both the B.1.1.7 virus and the ancestral virus Bavpat-1 (used in this study) contain the D614G mutation that was previously described to increase fitness in human airway cells and transmission between hamsters (*34, 40, 41*). One of the mutations present in B.1.1.7, but not in Bavpat-1, is an N501Y change in spike, which is an ACE2 contact residue, and has been proposed to increase the binding to human ACE2. This mutation is also present in the South-African (B.1.135) and Brazilian variant (P.1), pointing towards convergent evolution. A recent study used a reverse genetic approach to show that a virus containing only the N501Y transmitted more efficiently between hamsters (*42*). The P681H change in spike may play a role as well, as this mutation is located directly N-terminal from the RRARS MBCS, which was shown to be critical for serine protease-mediated entry into human airway cells (*11*). P681H adds an additional basic residue to the MBCS and increases S1/S2 cleavage (*39, 43*). Another relevant mutation could be a stop codon early in the B.1.1.7 ORF8. This protein was shown to be secreted, leading to activation of IL17 signaling by binding to IL17RA (*44*). In addition, ORF8 was shown to interact with TGF-β (*45*), a cytokine thought to play an important role in lung inflammation, fibrosis and edema (*46, 47*). Whether dysregulating IL17 or TGF-β signaling is beneficial for virus replication needs to be investigated, but a secreted factor that generates a pro-viral milieu could explain why B.1.1.7 did not require more time to outcompete Bavpat-1 at a 5:1 compared to a 1:1 Bavpat-1-to-B.1.1.7 ratio, especially if there are costs associated with ORF8 expression. These results support an important role for social interactions in determining viral fitness (*48*). The possibility that specific virus-host interactions may be crucial in determining the outcome of experiments that measure viral fitness, and such interactions can be lost in animal models. Future studies may use reverse genetics to address the roles of these mutations in human organoid systems.

An important question related to variants such as the B.1.1.7 clade is whether they are associated with increased health risks and mortality (*4, 5, 49*). In our study we did not observe major differences in the host response between B.1.1.7 and Bavpat-1-infected organoids suggesting that an altered host response is unlikely to drive potential differences in disease severity. Interestingly, Frampton and colleagues noted a trend towards a ∼2 day longer time to hospitalization from symptom onset for patients infected with B.1.1.7 (median 6 days) compared with patients infected with a different clade (median 4 days), which could reflect the observed replication kinetics on 2D AT2 cells (Fig. 4H-I) (*5*).

In conclusion, we report that extended shedding is an *in vitro* correlate of infectiousness for SARS-CoV-2 B.1.1.7, which corroborates epidemiological findings and may explain why this clade is more transmissible. Human adult stem cell-derived pulmonary and intestinal organoids thus may provide a platform for the comparison of non-cell culture-adapted SARS-CoV-2 clinical isolates.

## Acknowledgments

We thank Robbert Rottier for providing human adult lung material, and the Microscopy CORE lab at Maastricht University for the TEM sample preparation. This manuscript was part of the research programme of the Netherlands Centre for One Health (www.ncoh.nl).

## Author contributions

Conceptualization: M.M.L., T.I.B., A.Z.M., B.L.H.

Methodology: M.M.L., T.I.B., A.Z.M., Y.W., D.C.W., N.C.W.

Validation: M.M.L., T.I.B., A.Z.M., J.V., H.C., B.L.H.

Formal analysis: M.M.L., T.I.B., A.Z.M., J.Z., Y.W., D.C.W., N.C.W.

Investigation: M.M.L., T.I.B., A.Z.M., K.K., N.G., C.D.K., J.Z., P.B.D., T.B., Y.W., D.C.W., N.C.W.

Resources: C.H.G., H.C., N.C.W., M.M., M.M.L., B.L.H.

Writing - Original Draft: M.M.L., T.I.B., A.Z.M., B.L.H.

Writing - Review & Editing: M.M.L., T.I.B., A.Z.M., N.C.W, H.C., B.L.H.

Visualization: M.M.L., T.I.B., A.Z.M., Y.W., D.C.W., N.C.W.

Supervision: M.M.L, B.L.H.

Funding acquisition: B.L.H.

## Data and materials availability

Messenger RNA sequencing data will be deposited in GEO. The consensus sequence of the passage 3 B.1.1.7 virus (which was identical to the sequence of the clinical sample) was submitted to Genbank (Accession no. MW947280). Raw deep-sequencing data are available at the NIH Short Read Archive under accession number: BioProject PRJNA722947. All pipelines and scripts for deep-sequencing data analysis are available on GitHub: https://github.com/nicwulab/UK_strain_in_vitro_fitness

## Materials and Methods

### Cell lines

VeroE6 cells were maintained in Dulbecco’s modified Eagle’s medium (DMEM, Gibco) supplemented with 10% fetal bovine serum (FBS), HEPES (20 mM, Lonza), sodium pyruvate (1 mM, Gibco), penicillin (100 IU/mL), and streptomycin (100 IU/mL) at 37°C in a humidified CO_2_ incubator. Calu-3 cells were maintained in Opti-MEM I (1) + GlutaMAX (Gibco) supplemented with 10% FBS, penicillin (100 IU/mL), and streptomycin (100 IU/mL) at 37°C in a humidified CO_2_ incubator. Cell lines were tested negative for mycoplasma.

### Airway cell isolation, culture and differentiation

Human adult airway stem cells were isolated as described previously (*22*) using a protocol adapted from Sachs and colleagues (*19*). Cells were differentiated using two methods. The first method used commercially available Pneumacult-ALI medium (complete base medium with 1X maintenance supplement; Stemcell) as described before (*22*). The second method made use of a defined medium, in which cells were (50,000 for 6.5 mm inserts; 200,000 for 12 mm inserts) dissociated using TrypLE Express (Gibco) and plated on rat tail collagen type I (20 µg/ml; Corning) coated transwell inserts (Corning) in airway organoid medium (AO, (*19*)). After reaching confluency, the medium in the lower compartment was replaced with Advanced DMEM/F12 (Gibco), supplemented with HEPES, Glutamax, penicillin (100 IU/mL) and streptomycin (100 IU/mL) (AdDF+++) containing *N*-acetylcysteine (1.25 mM; Sigma), B27 (1X, Gibco) with or without BMP2 (50 ng/ml; Peprotec) while the medium in the upper compartment was removed to establish an air-liquid interface. Cells were differentiated at the air-liquid interface for 3-4 weeks. Medium was replaced every 4-5 days.

Adult human lung tissue for airway and alveolar stem cell (see below) isolation was obtained from non-tumour lung tissue obtained from patients undergoing lung resection. Lung tissue was obtained from residual, tumor-free, material obtained at lung resection surgery for lung cancer. The Medical Ethical Committee of the Erasmus MC Rotterdam granted permission for this study (METC 2012-512).

### Alveolar cell isolation, culture and differentiation

Human adult alveolar stem cells were isolated using a protocol adapted from Lamers and colleagues (2020). Distal lung pieces were chopped into 2 x 2 mm pieces, washed twice with cold PBS and incubated in 100% dispase (Corning), supplemented with 10 µM Y-27632 (MedChemExpress) for 30-60 min at 37°C. Next, cells were dissociated from the tissue by pipetting using a 5 ml serological pipet and subsequently using a P1000 micropipette. Cells were then strained using a 100 µm cell strainer (Falcon) and washed twice in cold AdDF+++. Red blood cells were lysed using red blood cell lysis buffer (Roche), after which cells were again washed twice with cold AdDF+++. Cells were pelleted, incubated on ice for 2 min, and plated in growth factor reduced basement membrane extract (BME; Cultrex). After BME solidification, alveolar medium (recipe (see below) adapted from Youk and colleagues (*27*)) was added and cells were incubated at 37°C in a humidified CO_2_ incubator (*27*). Alveolar medium consisted of AdDF+++ containing B27 (1X; minus vitamin A; Invitrogen), N2 (1X; Invitrogen), *N*-acetylcysteine (1.25 mM), CHIR99021 (3 µM; Sigma), RSPO1 (R-spondin 1; conditioned medium, 10% v/v), FGF7 (100 ng/ml; Peprotec), FGF10 (100 ng/ml; Peprotec), epidermal growth factor (EGF; 50 ng/ml; Peprotec), NOG (NOGGIN; 100 ng/ml; Peprotec), SB431542 (10 µM; Tocris), and Primocin (1X; Invivogen). Y-27632 (10 µM) was added for the first 5 days. Medium was replaced every 4-5 days.

To sort AT2 cells, organoids were digested using TrypLE Express and washed in AdDF++ twice before incubating the cells in Lysotracker (Thermofisher) for 20 min at 37°C. Next, cells were incubated in FACS buffer (2mM EDTA, 2.5% bovine serum albumin (BSA) in PBS) on ice for 5 min, stained with the AT2 marker antibody HTII-280 (1:40; Terrace Biotech) on ice for 15 min, and with goat anti-mouse IgM Alexa Fluor 488 (1:400; Invitrogen) for 5 min. Cells were then washed once in FACS buffer and HTII-280-high and Lysotracker-high cells were sorted using a FACS Aria cell sorter (BD biosciences) into AdDF+++ containing 10 µM Y-27632. Cells were pelleted, incubated on ice for 2 min and plated in BME domes in alveolar medium. Y-27632 (10 µM) was added for the first 5 days. Cells were incubated at 37°C in a humidified CO_2_ incubator and medium was replaced every 4-5 days.

Sorted and unsorted AT2 organoids were split every 2-3 weeks at a 1:3-1:8 ratio. Organoids were dissociated to small clumps using TrypLE.

For 2D experiments, sorted AT2 organoids were dissociated using TrypLE, washed in AdDF+++ twice, and plated on BME-coated transwell inserts in alveolar medium containing Y-27632 (10 µM). Inserts were coated with BME at a 1:100 dilution in PBS at 37°C for 30 min. Cells were incubated at 37°C in a humidified CO_2_ incubator for 4 days (until a confluency of ∼80% was reached), after which the medium was replaced with AdDF+++ supplemented with 10% FBS. In this medium, the cells grew to confluency within 3 days. Next, the medium in the top compartment was removed to establish an air-liquid interface and the medium in the bottom compartment was replaced for AdDF+++ without FBS. These cells were used for infections directly.

All centrifugation steps were performed at 400 xg for 3 min.

### Intestinal organoid isolation and culture

Human small intestinal cells were isolated and cultured as previously described (*50, 51*). Tissue from the ileum was obtained from the UMC Utrecht with informed consent of the patient, who was operated for resection of an intestinal tumor. A sample from non-transformed, normal mucosa was taken for this study. The study was approved by the UMC Utrecht (Utrecht, The Netherlands) ethical committee and was in accordance with the Declaration of Helsinki and according to Dutch law. This study is compliant with all relevant ethical regulations regarding research involving human participants.

For 2D experiments, human small intestinal organoids were mechanically disrupted, dissociated using TrypLE, washed twice in AdDF+++, and plated as single cells in human small intestinal medium supplemented with Y-27632 (10 µM) on wells coated with 1:100 BME and 20 µg/ml rat tail collagen type I. Monolayers were grown until 70-90% confluency and subsequently used for infections.

### Virus isolation and propagation

SARS-CoV-2 (isolate BetaCoV/Munich/BavPat1/2020; European Virus Archive Global #026V 03883; kindly provided by Dr. C. Drosten) and SARS-CoV-2 B.1.1.7 (Genbank MW947280) were propagated as described before (*10*). Briefly, the indicated viruses were propagated to passage 3 on Calu-3 cells in AdDdF+++ at 37°C in a humidified CO_2_ incubator. Infections were performed at a multiplicity of infection (moi) of 0.01 and virus was harvested after 72 hours or at the peak of replication. The culture supernatant was cleared by centrifugation, filtered by a 0.45 µM low protein binding filter (Millipore) and medium was exchanged three times for Opti-MEM I (1X) + GlutaMAX using an Amicon Ultra-15 column (100 kDa cutoff). Next, the supernatants were stored at −80°C. Virus titers were determined by plaque assay as described below. All work with infectious SARS-CoV-2 was performed in a Class II Biosafety Cabinet under BSL-3 conditions at Erasmus Medical Center.

### SARS-CoV-2 infections

Prior to infection, cells were washed three times using AdDF+++. Washes were performed apically for 2D cultures or in 15 ml tubes for 3D cultures with 300 µl or 5 ml medium, respectively. Cells were incubated with inoculums for 4 hours at 37°C in a humidified CO_2_ incubator at a moi of 0.1.

Washing was performed three times 4 hours post infection in AdDF+++ after which:

- For 2D ALI cultures, a fourth wash was performed and collected as the 4 hour post-infection baseline.
- For 2D submerged cultures, 500 µl expansion medium (for intestinal organoids) or OptiMEM I (1X) + GlutaMAX (for VeroE6 and Calu-3 cells) was added after three washes and a sample was taken from the supernatant as the 4 hour post-infection baseline.
- For 3D cultures, a fourth wash was performed in AdDF+++, after which organoids were plated in 30 µl BME droplets in 48 well plates. 300 µl expansion medium was added after BME solidification and a 4 hour post-infection baseline sample was taken.

For 2D ALI cultures, samples were taken at the indicated time points as follows: 500 µl for 12mm inserts or 200µl for 6.5mm inserts of AdDF+++ was added to the apical side of the cells and cells were incubated for 10 min at 37°C in a humidified CO_2_ incubator after which supernatants were pipetted up and down on the cells twice and transferred to a microvial. For 2D submerged cultures, 200µl samples were taken directly from the supernatant and replaced with 200 µl fresh medium. For 3D cultures, samples were taken by removing the supernatant and resuspending the organoids along with the BME in 300 µl AdDF+++. Organoid suspensions were transferred to a microvial before freezing to lyse the organoids. All samples were stored at - 80°C until further processing for viral titer determinations. At 3 days postinfection light microscopy images were taken of VeroE6 infected cultures using a Zeiss Primovert microscope and analysed using ZEN software (Zeiss).

### Determination of virus titers using qRT-PCR

SARS-CoV-2 RNA was extracted as described previously (*21*) and RNA genome copies (E-gene) were determined by qRT-PCR. Briefly, supernatant or organoid samples were thawed and centrifuged at 500 x g (3D organoids) or 2,000 x g (supernatant) for 3 min. 60 µl sample was lysed in 90 µl MagnaPure LC Lysis buffer (Roche) at RT for 10 min. RNA was extracted by incubating samples with 50 µl Agencourt AMPure XP beads (Beckman Coulter) for 15 min at room temperature, washing beads twice with 70% ethanol on a DynaMag-96 magnet (Invitrogen) and eluting in 30 µl DEPC-treated water. RNA copies per ml were determined by qRT-PCR using primers targeting the E gene (*52*) and comparing the Ct values to a counted standard curve derived from a Bavpat-1 stock titrated on VeroE6 cells. Where indicated beta actin was used a housekeeping gene using the following primers and probe: Bact_fw GGCATCCACGAAACTACCTT, Bact_rev AGCACTGTGTTGGCGTACAG Bact_probe FAM-ATCATGAAGTGTGACGTGGACATCCG-BHQ1.

### Viral titer determinations by plaque assay

Virus stocks or experimental samples were thawed and diluted in 10-fold serial dilutions in 200µl Opti-MEM I (1X) + GlutaMAX. 100µl of each dilution was added to monolayers of 1 × 10^6^ VeroE6 or Calu-3 cells in the same medium in a 12-well plate. Cells were incubated with inoculums at 37°C for 4 hours and then medium was replaced for 1.2% Avicel (FMC biopolymers) in Opti-MEM I (1X) + GlutaMAX for two days. Cells were fixed in 4% formalin for 20 minutes, permeabilized in 70% ice-cold ethanol and washed in PBS. Cells were blocked in 0.6% BSA (bovine serum albumin; Sigma) in PBS and stained with mouse anti-nucleocapsid (Sino biological; 1:2000) in PBS containing 0.1% BSA, washed thrice in PBS, and stained with goat anti-mouse Alexa Fluor 488 (Invitrogen; 1:4000) in PBS containing 0.1% BSA. Cells were washed thrice in PBS and plates were scanned on the Amersham Typhoon Biomolecular Imager (channel Cy2; resolution 10 µm; GE Healthcare). All staining steps were performed at room temperature for one hour. Plaque assay analysis was performed using ImageQuant TL 8.2 software (GE Healthcare).

### Immunofluorescent staining

Transwell inserts and 2D intestinal cultures were fixed in 4% formalin for 20 min, permeabilized in 70% ethanol for airway and intestinal cells and 0.1% Triton X-100 (Sigma) for alveolar cells, and blocked for 60 min in blocking buffer (10% filtered normal goat serum (MP Biomedicals) in PBS. Cells were incubated with primary antibodies overnight at 4°C in blocking buffer, washed twice with PBS, incubated with corresponding secondary antibodies (Alexa 488-/ 594-conjugated anti-rabbit IgG, anti-mouse IgG, or anti-mouse IgG1 (1:500; Invitrogen)) in blocking buffer for 2 hours at room temperature, washed two times with PBS, incubated with indicated additional stains (hoechst (ThermoFisher) or phalloidin (Santa Cruz)), washed twice with PBS, and mounted in Prolong Antifade (Invitrogen) mounting medium. Viral nucleocapsid was stained with rabbit anti-NP (1:500; Sino biological), ciliated cells were stained with anti-AcTub (1:100; Santa Cruz, 6-11B-1), club cells with anti-CC10 (1:200; Santa Cruz,), basal cells with anti-TP63 (1:200, Abcam), goblet cells with anti-MUC5AC (1:200, Thermofisher), TMPRSS2 with anti-TMPRSS2 (1:400, Abcam) and AT2 cells with anti-HTII-280 (1:200, Terrace Biotech).

For ACE2 stainings, cells were blocked with 1% BSA and stained overnight with goat anti-ACE2 (1:200, R&D Systems) in blocking buffer followed by rabbit anti-goat Alexa Fluor 488 (1:500, Invitrogen) in blocking buffer for 2 hours at room temperature, washed two times with PBS, incubated with indicated additional stains (hoechst or phalloidin), washed twice with PBS, and mounted in Prolong Antifade mounting medium

3D Alveolar and airway organoids were fixed in 4% formalin overnight, permeabilized in 0.1% Triton X-100, and blocked for 60 min in 10% normal goat serum in PBS (blocking buffer). Cells were incubated with primary antibodies overnight at 4°C in blocking buffer, washed twice with PBS, incubated with corresponding secondary antibodies (Alexa488- and 594-conjugated anti-rabbit IgG, anti-mouse IgG, or anti-mouse IgG1 (1:500; Invitrogen)) in blocking buffer for 2 hours at room temperature, washed two times with PBS, incubated with indicated additional stains (hoechst or phalloidin), washed twice with PBS, and mounted in Prolong Antifade mounting medium. Viral nucleocapsid was stained with rabbit anti-NP, AT2 cells with anti-HTII-280 or anti-SFPTC (Millipore) and basal cells with anti-TP63.

All confocal imaging was performed on a LSM700 confocal microscope using ZEN software.

### Replicative fitness measurements

Replicative fitness was measured as described before (*34*). Briefly, air-liquid airway organoids differentiated in Pneumacult were infected as described above at a 1:1 or 5:1 Bavpat-1-to-B.1.1.7 ratio at a final moi of 0.1 (based of PFU/ml). Apical washes were performed daily and stored at -80°C until further processing. RNA was extracted as described previously (*21*) and samples were subjected to cDNA synthesis using Superscript IV (Invitrogen), according to the manufacturer’s instructions. Next, a ∼700 bp region in the S gene containing the P681H mutation and a ∼300 bp NSP3 covering the 3675-3677 deletion were amplified by PCR using Pfu UltraII (Agilent) using the following primers:

P681H: ‘5-TGA CAC TAC TGA TGC TGT CCG TG-3’ (forward) and ‘5-GAT GGA TCT GGT AAT ATT TGT GG-3’ (reverse).

3675-3677 deletion: ‘5-ATG GGT ATT ATT GCT ATG TCT GC -3’ (forward) and ‘5-GTG TCA AGA CAT TCA TAA GTG TCC -3’ (reverse).

Conditions for the P681H PCR were as follows:

1. Denaturation for 5 minutes at 94°C;
2. Denaturation for 20 seconds at 94°C;
3. Annealing for 20 seconds at 52°C;
4. Extension for 60 seconds at 72°C;
5. Steps 2-4 40 times;
6. Final extension for 5 minutes at 72°C.

Conditions for the <u>3675-3677</u> deletion PCR were as follows:

1. Denaturation for 5 minutes at 94°C;
2. Denaturation for 20 seconds at 94°C;
3. Annealing for 20 seconds at 50°C;
4. Extension for 30 seconds at 72°C;
5. Steps 2-4 40 times;
6. Final extension for 5 minutes at 72°C.

Next, amplified PCR products were run on agarose gel to verify that the PCR worked and the remaining PCR products were purified using a PCR purification kit (Qiagen) before Sanger sequencing using the forward primer. Sanger sequences were analysed using QSVanalyser (Insilicase) or Synthego ICE CRISPR indel frequency analysis software. Raw peak heights were calculated at the second nucleotide of 681 codon of spike (cytosine (Bavpat-1) or adenine (B.1.1.7)), or at the first nucleotide (thymine (Bavpat-1) or adenine (B.1.1.7)) of the 3675-3677 NSP3 deletion using QSVanalyser. Replicative fitness was calculated according to w = (f0/i0), in which i0 is the initial Bavpat-1/B.1.1.7 ratio and f0 is the Bavpat-1/B.1.1.7 ratio at a given timepoint (*34*).

For the analysis using Synthego ICE, sanger sequences were uploaded and analysed using a Bavpat-1 control sequence as reference and TTGATACTAGTTTG as a guide to generate allele percentages of B.1.1.7 (3675-3677 del)) and Bavpat-1 (no deletion) present in the sample. Next, allele percentages were used to calculate the relative replicative fitness as described above. The lower limit of detection for the Synthego ICE analysis was set at 1% allele frequency.

### Illumina library preparation

Illumina library preparation was performed as described before (*10*). Briefly, RNA was extracted as for qRT-PCR. Subsequently, cDNA was generated using ProtoscriptII reverse transcriptase enzyme (New England, BiotechnologyBioLabs) according to the manufacturer’s protocol. Samples were either amplified using an in-house-developed SARS-CoV-2-specfic multiplex PCR (*53*) (for virus stocks produced on cell lines), or using the QIAseq SARS-CoV-2 Primer Panel (Qiagen) (for organoid supernatants). Amplicons were purified with 0.8x AMPure XP beads and 100ng of DNA was converted to paired-end Illumina sequencing libraries using the KAPA HyperPlus library preparation kit (Roche) with the KAPA unique dual-indexed adapters (Roche) as per manufacturers recommendations. The barcode-labeled samples were pooled and anlyzed on an Illumina sequencer V3 MiSeq flowcell (2×300 cycles).

### Data analysis of SARS-CoV-2 deep-sequencing

Analysis of SARS-CoV-2 genome sequencing was performed as described previously (*10*). Briefly, sequencing adapters from the paired-end sequencing reads were trimmed by cutadapt (*54*), and aligned to the corresponding genome (Bavpat-1 and B.1.1.7 isolate consensus sequence) using Bowtie2 (*55*) with the parameters: *--no-discordant --dovetail --no-mixed -- maxins 2000*. The first 30 nt from both 5’ and 3’ ends of all reads were soft-clipped using bamUtils (*56*) using parameters: -L 30 -R 0 --clip, to remove artifactual bases from SARS-CoV-2 primers that may interfere with variant calling. Variant calling was performed by VarScan2 (*57*) with parameters: pileup2cns --min-coverage 10 --min-reads2 2 --min-var-freq 0.01 --min-freq-for-hom 0.75 --p-value 0.05 --variants 1 using pileup data from SAMtools mpileup (*58*) with parameters: --excl-flags 2048 --excl-flags 256 --max-depth 50000 --min-MQ 30 --min-BQ 30. The sequence of the passage 3 B.1.1.7 virus was submitted to Genbank (MW947280). For the competition experiment, read mapping and variant calling were performed using Bavpat-1 as the reference sequence. The frequency of the B.1.1.7 strain in the sample was computed as the average frequency of all mutations that represent the difference between Bavpat-1 and the B.1.1.7 strain. Specifically, sequence logos were generated by logomaker (*59*). Raw sequencing data are available at the NIH Short Read Archive under accession number: BioProject PRJNA722947. All pipelines and scripts for data analysis are available on GitHub: https://github.com/nicwulab/UK_strain_in_vitro_fitness

### Bulk mRNA sequencing

Library preparation was performed at Single Cell Discoveries (Utrecht, The Netherlands), using an adapted version of the CEL-seq protocol. In brief: total RNA was extracted using the standard TRIzol (Invitrogen) protocol and used for library preparation and sequencing. mRNA was processed as described previously, following an adapted version of the single-cell mRNA seq protocol of CEL-Seq (*60, 61*). Samples were barcoded with CEL-seq primers during a reverse transcription and pooled after second strand synthesis. The resulting cDNA was amplified with an overnight *in vitro* transcription reaction. From this amplified RNA, sequencing libraries were prepared with Illumina Truseq small RNA primers. Paired-end sequencing was performed on the Illumina NextSeq500 platform using barcoded 1 x 75 nucleotide read setup. Read 1 was used to identify the Illumina library index and CEL-Seq sample barcode. Read 2 was aligned to the CRCh38 human RefSeq transcriptome, with the addition of SARS-CoV-2 (Ref-SKU: 026V-03883; MW947280) genomes, using BWA using standard settings (*62*). Reads that mapped equally well to multiple locations were discarded. Mapping and generation of count tables was done using the MapAndGo script (https://github.com/anna-alemany/transcriptomics/tree/master/mapandgo). Data will be deposited in GEO. SARS-CoV-2-mapping reads were removed for further analysis. Differential expression analysis were performed using the DESeq2 package (*63*). P-values resulting from the differential expression tests were adjusted with Benjamini & Hochberg correction and considered significant with Padj<0.05. Shrinkage is applied to the resulting fold changes. To illustrate the results, the significantly differentially regulated genes with an absolute fold change above 2 (with a maximum of 50 genes) are depicted in heatmaps, using expression values after variance stabilizing transformation.

### Transmission electron microscopy

Organoids were chemically fixed for 3 hours at room temperature with 1.5% glutaraldehyde in 0.067 M cacodylate (pH 7.4) and 1 % sucrose. Samples were washed once with 0.1 M cacodylate (pH 7.4), 1 % sucrose and 3 times with 0.1 M cacodylate (pH 7.4), followed by incubation in 1% osmium tetroxide and 1.5% K4Fe(CN)6 in 0.1 M sodium cacodylate (pH 7.4) for 1 hour at 4°C. After rinsing with Milli-Q water, organoids were dehydrated at room temperature in a graded ethanol series (70%, 90%, up to 100%) and embedded in epon. Epon was polymerized for 48h at 60°C. Ultrathin sections (60 nm) were cut using a diamond knife (Diatome) on a Leica UC7 ultramicrotome, and transferred onto 50 Mesh copper grids covered with a formvar and carbon film. Sections were post-stained with uranyl acetate and lead citrate. All TEM data were collected autonomously as virtual nanoscopy slides (*64*) on FEI Tecnai T12 microscopes at 120kV using an Eagle camera. Data was stitched, uploaded, shared and annotated using Omero (annotated using Omero) and PathViewer.

### Statistical analysis

Statistical analysis was performed with the GraphPad Prism 9 software. We compared differences in virus titers by a two-way ANOVA followed by a Sidak multiple-comparison test or by an unpaired T-test, as indicated in the figure legends.

**Supplementary table 1.** Genetic characterization of the isolated SARS-CoV-2 B.1.1.7 compared with the Bavpat-1 reference.

---

**Supplementary figure 1.**
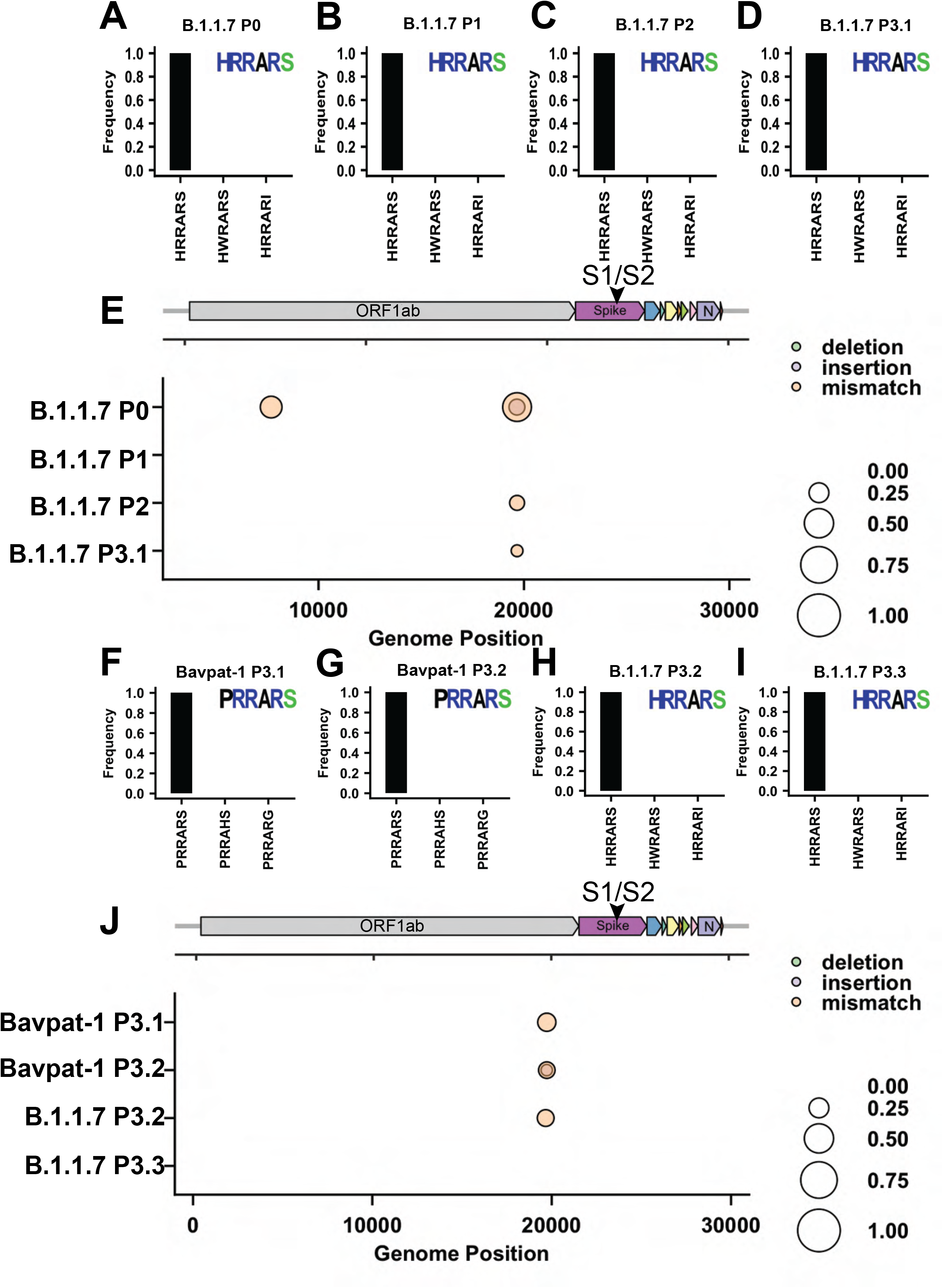
Deep-sequencing genetic characterization of the Calu-3 SARS-CoV-2 B.1.1.7 isolate. (A-D) Frequency plots of multibasic cleavage site (MBCS) mutations of the clinical sample (P0; A) and clinical isolates at passages 1-3 (P1-3; B-D). (E) Full genome frequency plots of P0-P3. (F-I) Frequency plots of MBCS mutations of the Bavpat-1 Calu-3 P3 stocks (replicate 1 = F; replicate 2 = G) and two B.1.1.7 Calu-3 P3 stocks (replicate 2 = H; replicate 3 = I) produced simultaneously. (J) Full genome frequency plots of Bavpat-1 (replicate 1 and 2) and B.1.1.7 (replicate 2 and 3) P3 stocks. MBCS plots contain the consensus MBCS amino acid sequence logo. In all plots variants with a frequency >10% are depicted.

**Supplementary figure 2.**
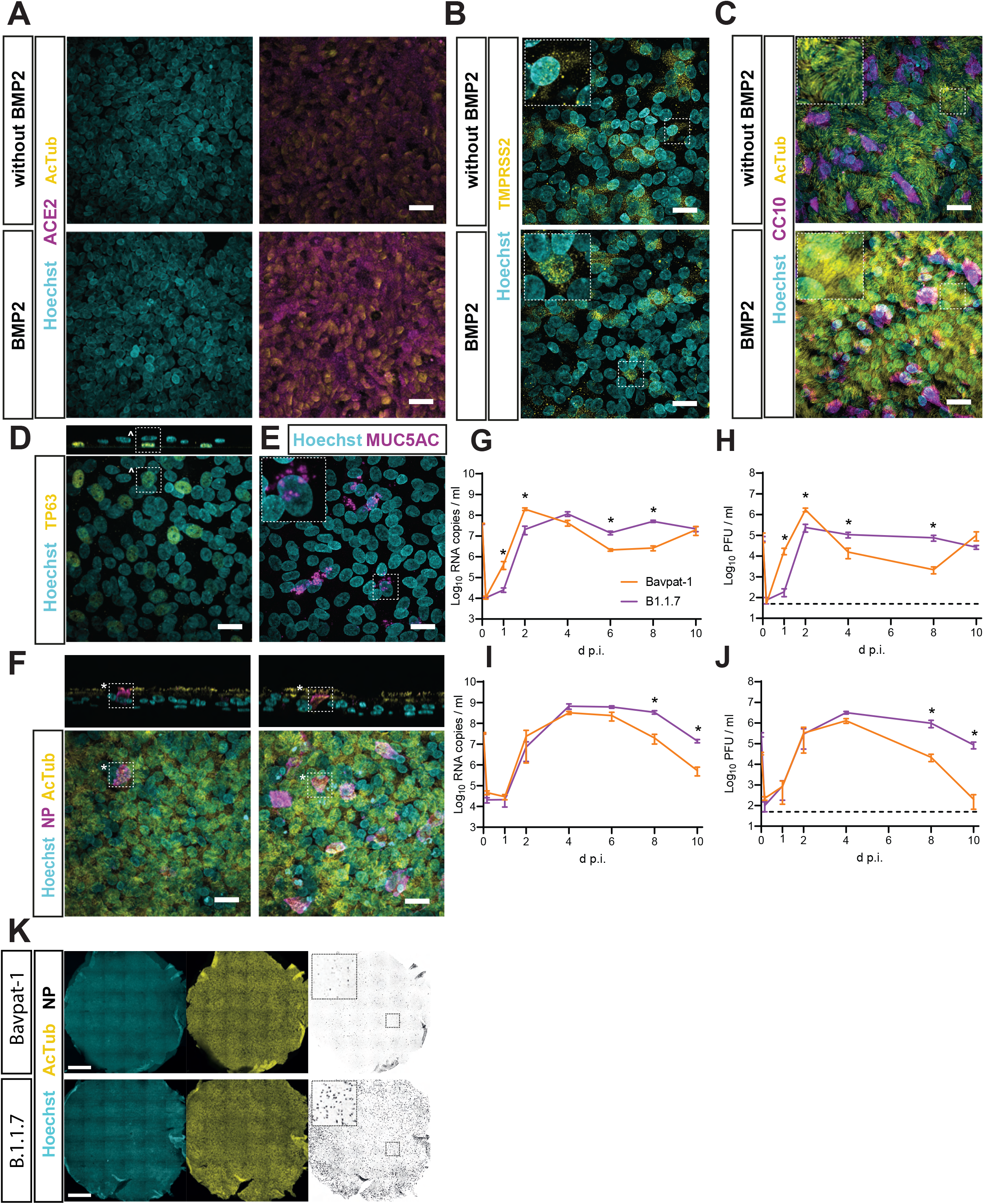
2D airway organoids grown at air-liquid interface in a defined medium containing BMP2 are well-differentiated and shed higher levels of SARS-CoV-2 B.1.1.7 compared with Bavpat-1 at late timepoints. (A-C) Immunofluorescent staining of 2D airway organoids differentiated with or without BMP2 for ACE2 (magenta; A), TMPRSS2 (yellow; B) and the ciliated cell marker acetylated tubulin (yellow, AcTub; C) alongside the club cell marker SCGB1A1 (magenta). (D-E) Immunofluorescent staining of BMP-2 differentiated 2D airway organoids for basal (yellow, TP63; D) and goblet (magenta, MUC5AC; E) cell markers. TP63 positive cells were only located basally (^). (F) Immunofluorescent staining of viral nucleoprotein (NP, magenta) and AcTub (yellow) showing infection of ciliated cells (*) by both B.1.1.7 and Bavpat-1. Scale bars in A to F indicate 20 µm. (G-J) Replication kinetics of B.1.1.7 and Bavpat-1 on BMP2 differentiated airway organoids from two donors (donor 1, G-H; donor 2, I-J). G and I show viral RNA copies (E gene) quantified by qPCR. H and J show infectious virus titers quantified by plaque assay on Calu-3 cells. (K) Full well images of Bavpat-1 and B.1.1.7 infected 2D airway organoids at day 10 post-infection. Nuclei are stained with hoechst (cyan). Scale bar indicates 1 mm. Dotted lines indicate detection limit (50 PFU/ml). D p.i. = days post-infection. Statistical analysis was performed by two-way ANOVA. * indicates a significant difference (P<0.05). Error bars indicate SEM. N=4 in G-J.

**Supplementary figure 3.**
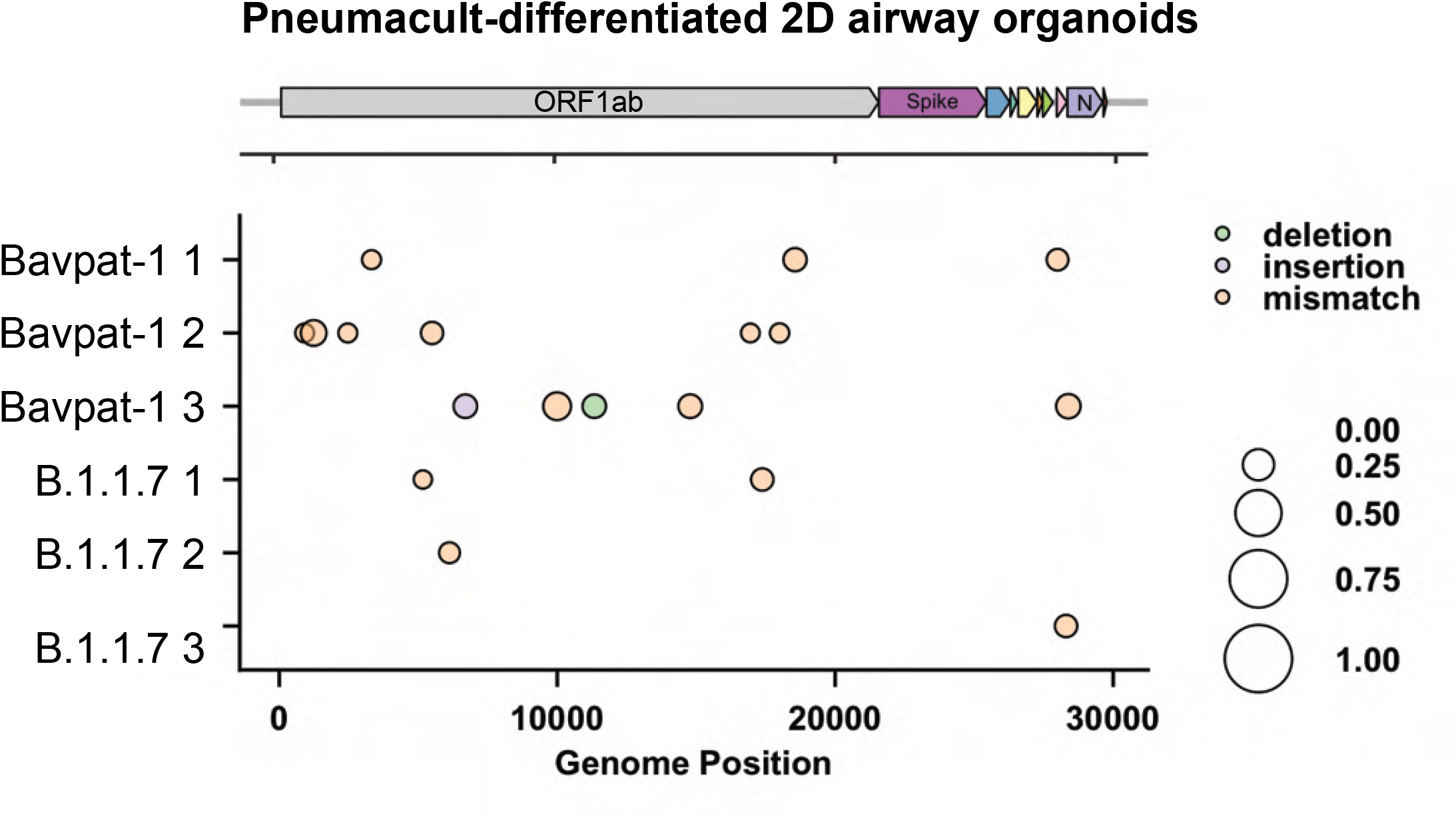
SARS-CoV-2 B.1.1.7 and Bavpat-1 are genetically stable on Pneumacult differentiated 2D human airway organoids. Full genome frequency plots of virus variants in apical washes of pneumacult-differentiated 2D airway organoids at day 10 post-infection for both the Bavpat-1 and B.1.1.7 isolate (three replicates each). In all plots variants with a frequency >10% are depicted.

**Supplementary figure 4.**
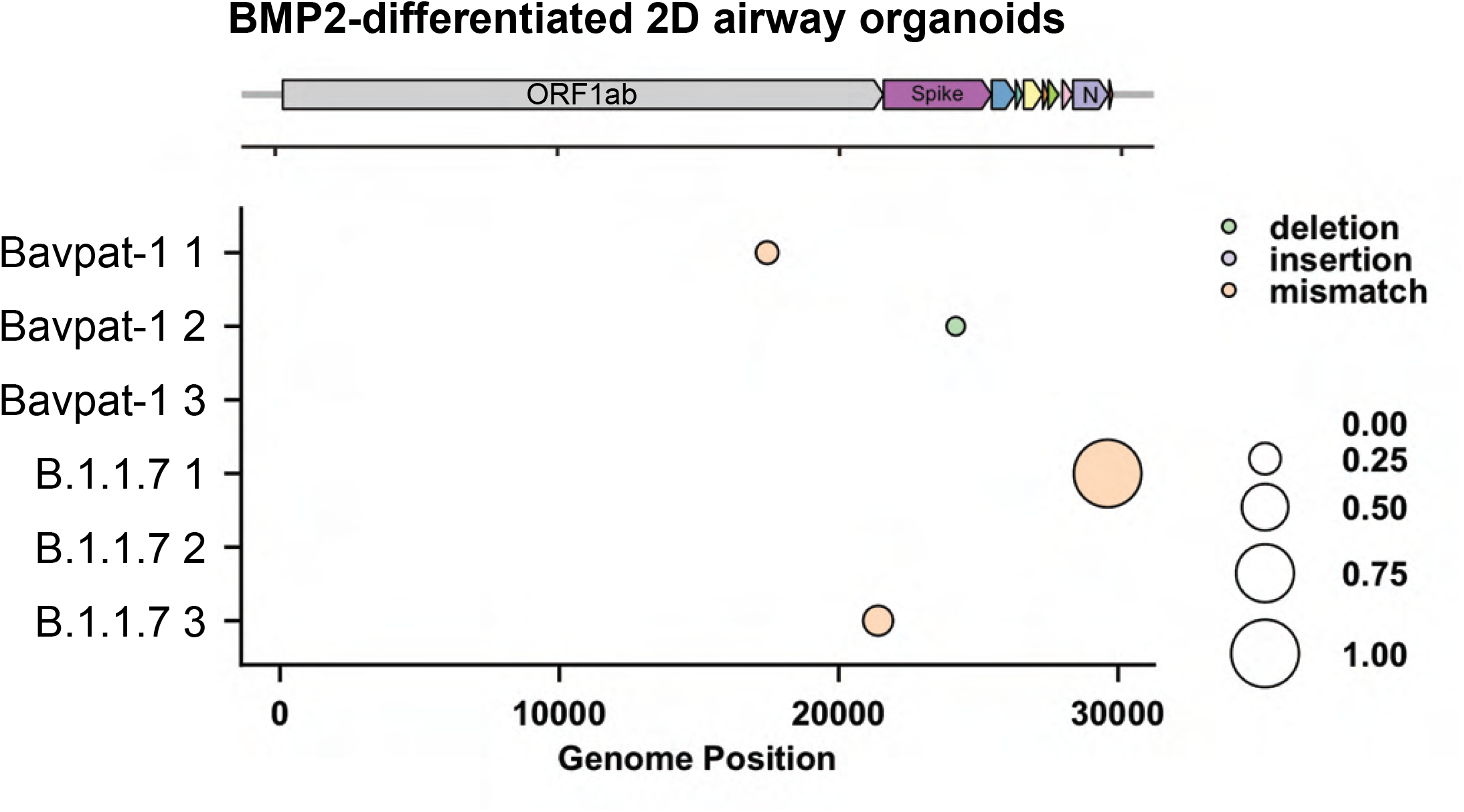
SARS-CoV-2 B.1.1.7 and Bavpat-1 are genetically stable on BMP2 differentiated 2D human airway organoids. Full genome frequency plots of virus variants in apical washes of BMP2-differentiated 2D airway organoids at day 10 post-infection for both the Bavpat-1 and B.1.1.7 isolate (three replicates each). In all plots variants with a frequency >10% are depicted.

**Supplementary figure 5.**
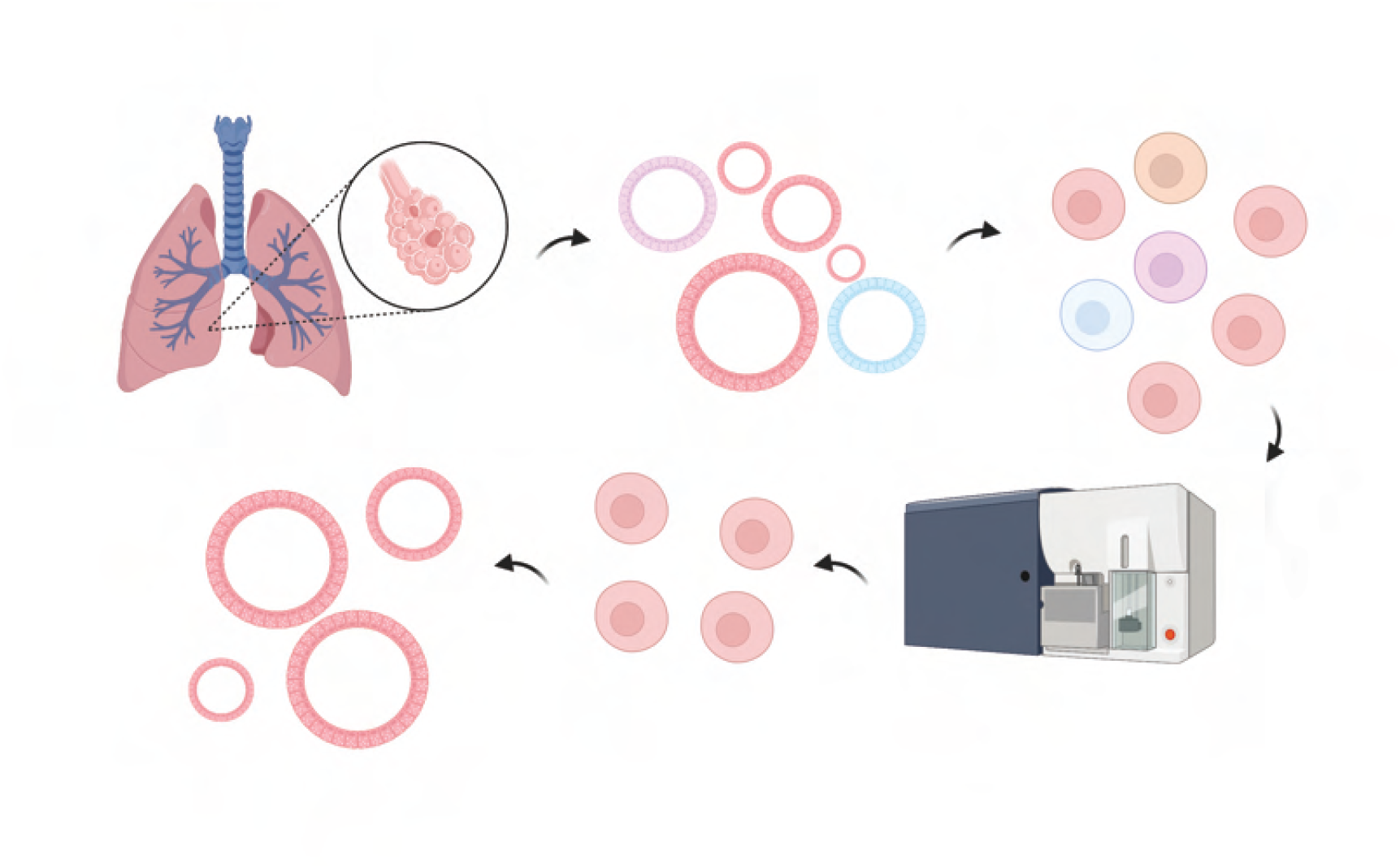
Schematic overview of the generation of human adult alveolar type 2 (AT2) organoids. Human adult stem cells are isolated from healthy lung tissue after lung resection surgery by enzymatic digestion of the tissue. Isolated stem cells are grown for 1 passage in basement membrane extract before they are dissociated and sorted for HTII-280 and Lysotracker high positivity to isolate a pure population of AT2 cells, which are again grown in basement membrane extract to generate AT2 organoids. Image created with BioRender.com

**Supplementary figure 6.**
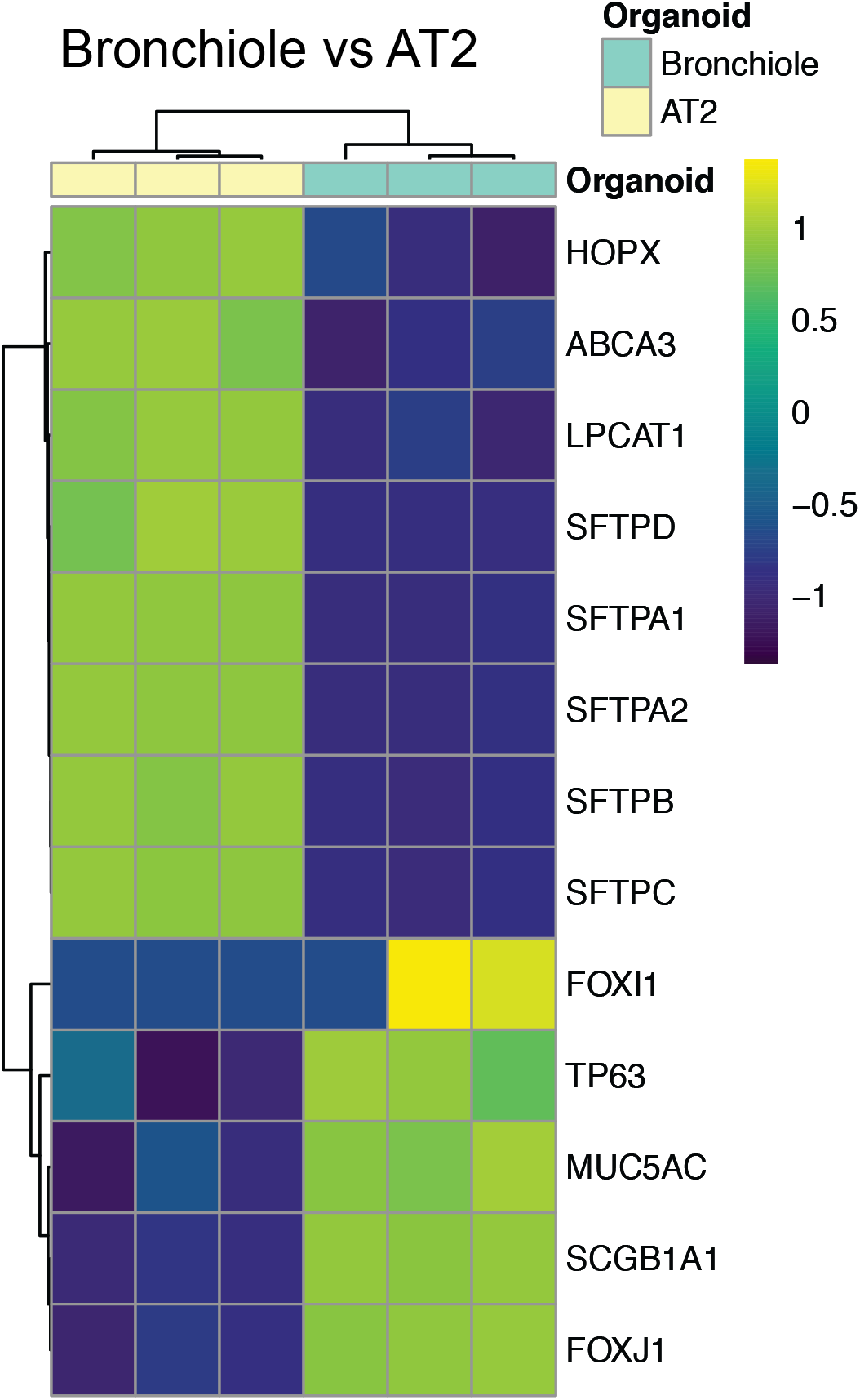
Human alveolar type 2 (AT2) organoids express AT2 - but not airway - marker genes. (A) Heatmap of normalized expression values scaled across samples of selected airway and alveolar marker genes in bronchiole and AT2 organoids. HOPX, ABCA3, LPCAT1, SFTPD, SFTPA1, SFTPA2, SFTPB and SFTPC are AT2 markers. FOXI1, TP63, MUC5AC, SCGB1A1 and FOXJ1 are markers for ionocytes, basal cells, goblet cells, club cells and ciliated cells, respectively.

**Supplementary figure 7.**
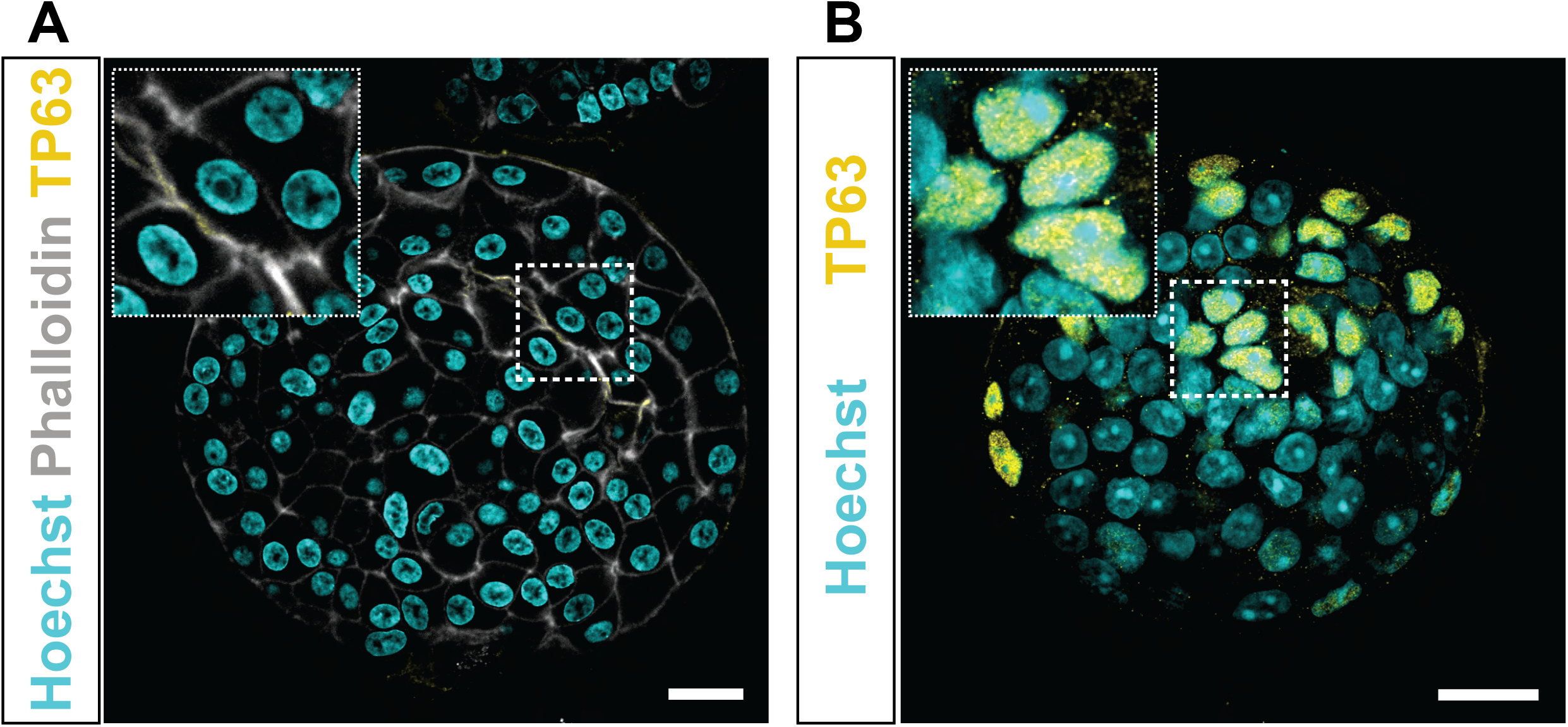
Human adult alveolar type 2 (AT2) organoids do not express the basal cell marker TP63. (A-B) Immunofluorescent staining of AT2 organoids (A) and bronchiolar organoids (B) for the basal cell marker TP63 (yellow). Nuclei are stained with hoechst (cyan) and actin is stained with phalloidin (white). Scale bars indicate 20 µm.

**Supplementary figure 8.**
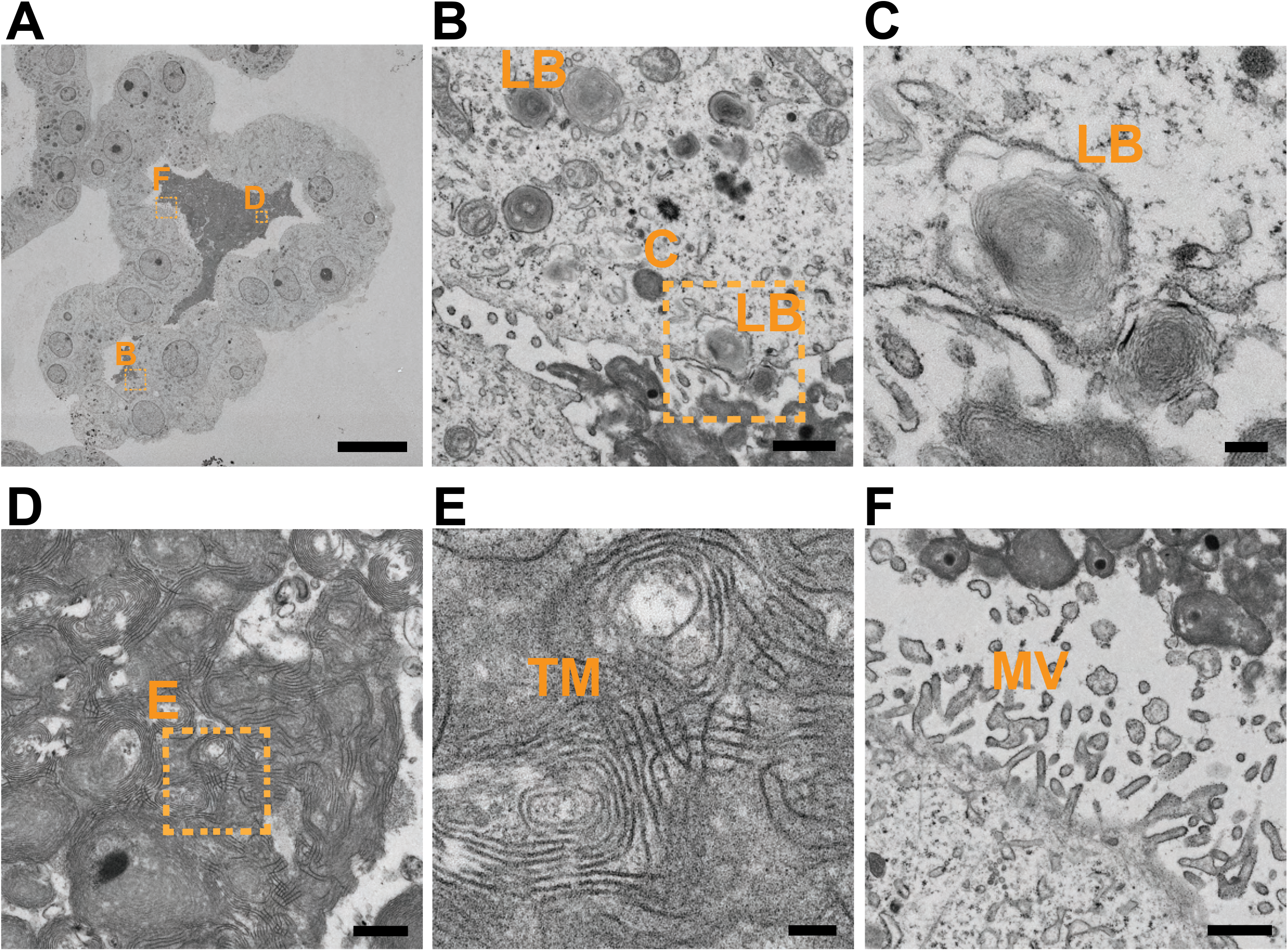
Transmission electron microscopy analysis of human adult alveolar type 2 organoids. (A-F) Overview of an intact alveolar type 2 (AT2) organoid showing cytoplasmic lamellar bodies (LB) (B and C) and secreted lamellar bodies (D and E) that organize into tubular myelin (TM). (F) Apical microvilli (MV) are predominant on the AT2 cells. Scale bars indicate 20 µm (A), 1 µm (B and F), 200 nm (C), 500 nm (D) and 100 nm (E).

**Supplementary figure 9.**
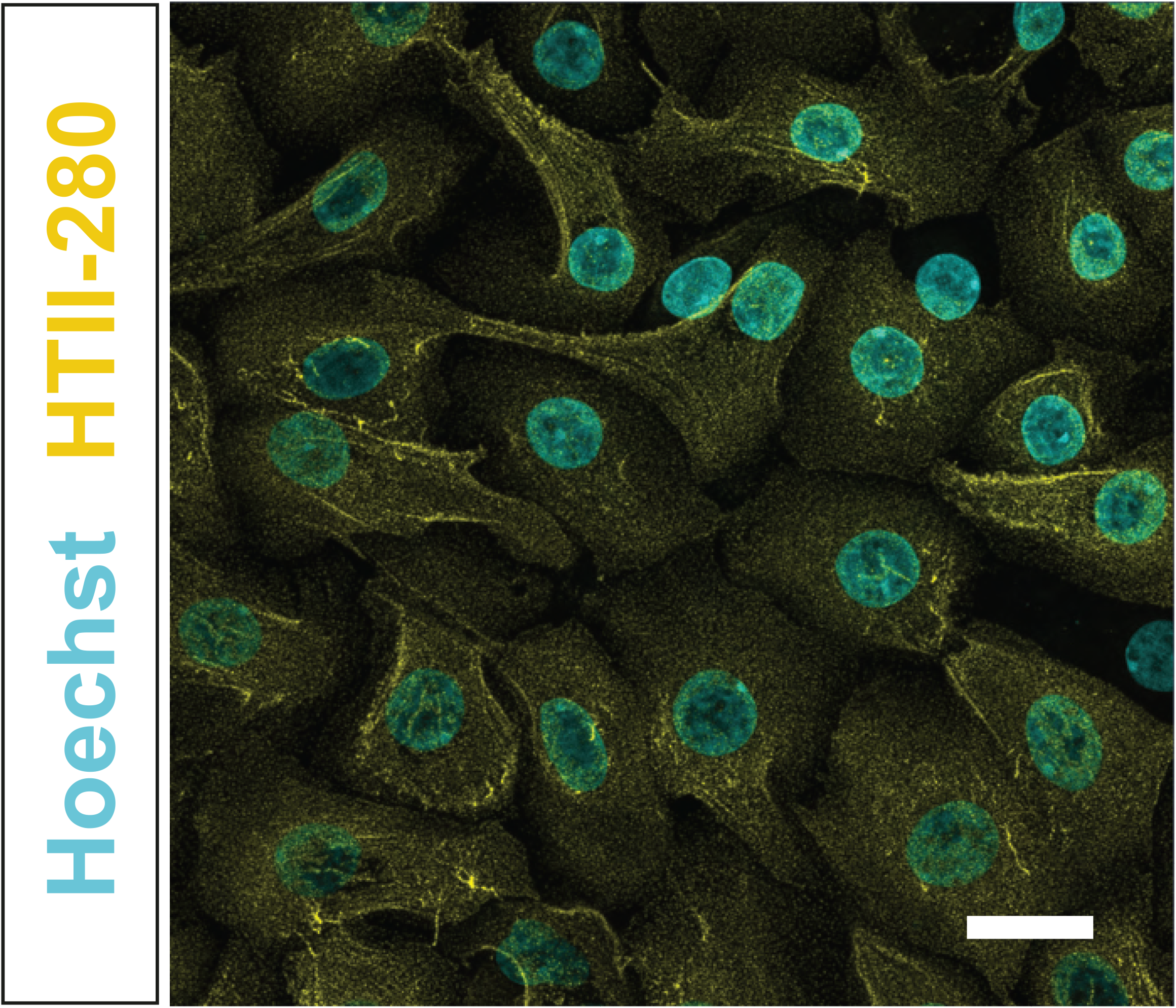
2D plated alveolar type 2 (AT2) cells stained for HTII-280. Immunofluorescent staining of 2D AT2 cells using HTII-280 (yellow). Nuclei are stained with hoechst (cyan). Scale bar depicts 20 µm.

**Supplementary figure 10.**
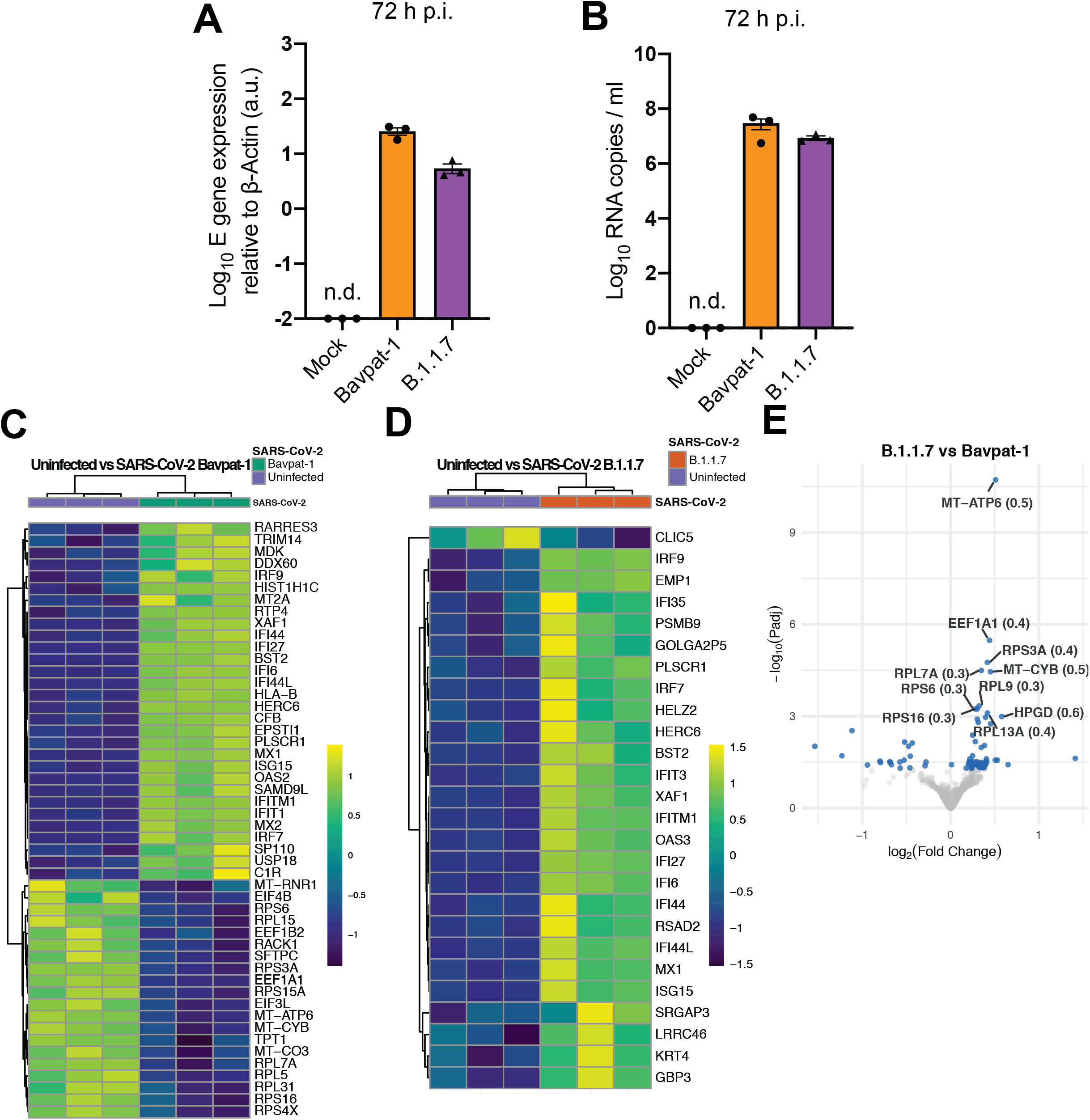
Host mRNA responses to SARS-CoV-2 B1.1.7 and Bavpat-1 infection in alveolar type 2 (AT2) organoids. (A-B) E gene expression (A) and viral RNA copies (B) in AT2 organoids infected with Bavpat-1 and B.1.1.7 at 72 h p.i.. (C-D) Heatmap of differentially expressed genes in response to Bavpat-1 (C) or B.1.1.7 (D) infection. Heatmaps show normalized expression of genes across samples. (E) Volcano plot depicting differentially expressed genes between Bavpat-1 and B.1.1.7-infected AT2 organoids. Blue dots indicate Padj<0.05. The names of the top 10 genes with the lowest Padj are shown. Error bars indicate SEM.

**Supplementary figure 11.**
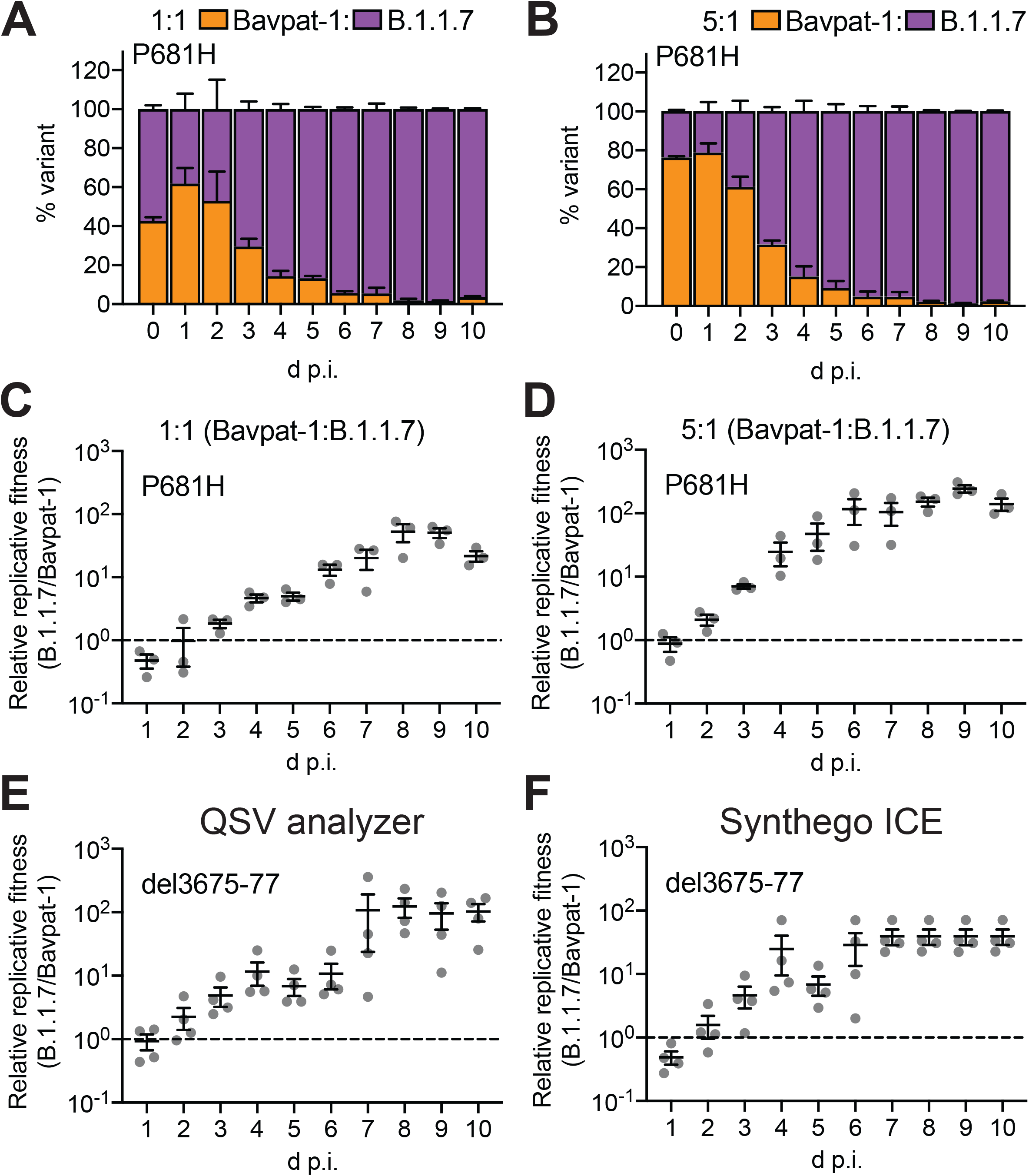
Confirmation of competition assay data. (A-B) Relative frequencies of Bavpat-1 and B.1.1.7 RNAs were assessed by RT–PCR and Sanger sequencing targeting the P681H mutation starting with an initial inoculum ratio of 1:1 (A) or 5:1 (B) (Bavpat-1:B.1.1.7), and a cumulative multiplicity of infection (moi) of 0.1 in airway donor 2. (C-D) These relative frequencies were used to calculate the relative replicative fitness at a 1:1 (C) and 5:1 (Bavpat-1:B.1.1.7) ratio (D). (E-F) Relative replicative fitness of Bavpat-1 and B.1.1.7 RNAs was assessed by RT–PCR and Sanger sequencing targeting the 3675-77 deletion starting with an initial inoculum ratio of 1:1, and a cumulative moi of 0.1 in airway donor 1. Sanger sequences were analysed with either QSVanalyser (Insilicase) (E) or Synthego ICE CRISPR indel frequency analysis software (F). The data in E and F was generated from the same competition experiment as Figure 6E and F. D p.i. = days post infection. Error bars indicate SEM.

## References

1. J. A. Plante et al., The variant gambit: COVID-19’s next move. Cell Host Microbe 29, 508–515 (2021).

2. S. Elbe, G. Buckland-Merrett, Data, disease and diplomacy: GISAID’s innovative contribution to global health. Glob Chall 1, 33–46 (2017).

3. K. Leung, M. H. Shum, G. M. Leung, T. T. Lam, J. T. Wu, Early transmissibility assessment of the N501Y mutant strains of SARS-CoV-2 in the United Kingdom, October to November 2020. Euro Surveill 26, (2021).

4. M. S. Graham et al., Changes in symptomatology, reinfection, and transmissibility associated with the SARS-CoV-2 variant B.1.1.7: an ecological study. Lancet Public Health, (2021).

5. D. Frampton et al., Genomic characteristics and clinical effect of the emergent SARS-CoV-2 B.1.1.7 lineage in London, UK: a whole-genome sequencing and hospital-based cohort study. Lancet Infect Dis, (2021).

6. E. Volz et al., Assessing transmissibility of SARS-CoV-2 lineage B.1.1.7 in England. Nature, (2021).

7. M. Richard et al., SARS-CoV-2 is transmitted via contact and via the air between ferrets. Nat Commun 11, 3496 (2020).

8. X. Montagutelli et al., The B1.351 and P.1 variants extend SARS-CoV-2 host range to mice. bioRxiv, 2021.2003.2018.436013 (2021).

9. W. B. Klimstra et al., SARS-CoV-2 growth, furin-cleavage-site adaptation and neutralization using serum from acutely infected, hospitalized COVID-19 patients. bioRxiv, (2020).

10. M. M. Lamers et al., Human airway cells prevent SARS-CoV-2 multibasic cleavage site cell culture adaptation. Elife 10, (2021).

11. A. Z. Mykytyn et al., SARS-CoV-2 entry into human airway organoids is serine protease-mediated and facilitated by the multibasic cleavage site. Elife 10, (2021).

12. S. Y. Lau et al., Attenuated SARS-CoV-2 variants with deletions at the S1/S2 junction. Emerg Microbes Infect 9, 837–842 (2020).

13. N. S. Ogando et al., SARS-coronavirus-2 replication in Vero E6 cells: replication kinetics, rapid adaptation and cytopathology. J Gen Virol, (2020).

14. H. Clevers, Modeling Development and Disease with Organoids. Cell 165, 1586–1597 (2016).

15. M. A. Lancaster, J. A. Knoblich, Organogenesis in a dish: modeling development and disease using organoid technologies. Science 345, 1247125 (2014).

16. K. Ettayebi et al., Replication of human noroviruses in stem cell-derived human enteroids. Science 353, 1387–1393 (2016).

17. X. Qian, H. N. Nguyen, F. Jacob, H. Song, G. L. Ming, Using brain organoids to understand Zika virus-induced microcephaly. Development 144, 952–957 (2017).

18. M. Watanabe et al., Self-Organized Cerebral Organoids with Human-Specific Features Predict Effective Drugs to Combat Zika Virus Infection. Cell Rep 21, 517–532 (2017).

19. N. Sachs et al., Long-term expanding human airway organoids for disease modeling. EMBO J 38, (2019).

20. J. Zhou et al., Differentiated human airway organoids to assess infectivity of emerging influenza virus. Proc Natl Acad Sci U S A 115, 6822–6827 (2018).

21. M. M. Lamers et al., SARS-CoV-2 productively infects human gut enterocytes. Science 369, 50–54 (2020).

22. M. M. Lamers et al., An organoid-derived bronchioalveolar model for SARS-CoV-2 infection of human alveolar type II-like cells. EMBO J, e105912 (2020).

23. J. Zhou et al., Infection of bat and human intestinal organoids by SARS-CoV-2. Nat Med 26, 1077–1083 (2020).

24. H. Katsura et al., Human Lung Stem Cell-Based Alveolospheres Provide Insights into SARS-CoV-2-Mediated Interferon Responses and Pneumocyte Dysfunction. Cell Stem Cell, (2020).

25. R. Pei et al., Host metabolism dysregulation and cell tropism identification in human airway and alveolar organoids upon SARS-CoV-2 infection. Protein Cell, (2020).

26. A. A. Salahudeen et al., Progenitor identification and SARS-CoV-2 infection in human distal lung organoids. Nature, (2020).

27. J. Youk et al., Three-Dimensional Human Alveolar Stem Cell Culture Models Reveal Infection Response to SARS-CoV-2. Cell Stem Cell, (2020).

28. J. J. A. van Kampen et al., Duration and key determinants of infectious virus shedding in hospitalized patients with coronavirus disease-2019 (COVID-19). Nat Commun 12, 267 (2021).

29. J. Van der Vaart, Clevers, H., Unpublished. (2021).

30. L. Zhao, M. Yee, M. A. O’Reilly, Transdifferentiation of alveolar epithelial type II to type I cells is controlled by opposing TGF-beta and BMP signaling. Am J Physiol-Lung C 305, L409–L418 (2013).

31. Y. Chen et al., The presence of SARS-CoV-2 RNA in the feces of COVID-19 patients. J Med Virol 92, 833–840 (2020).

32. W. Wang et al., Detection of SARS-CoV-2 in Different Types of Clinical Specimens. JAMA 323, 1843–1844 (2020).

33. C. C. Yong Zhang, Shuangli Zhu, Chang Shu, Dongyan Wang, Jingdong Song, Yang Song, Wei Zhen, Zijian Feng, Guizhen Wu, Jun Xu, Wenbo Xu., China CDC Weekly, 2020, 2(8): 123–124. doi: 10.46234/ccdcw2020.033, (Isolation of 2019-nCoV from a Stool Specimen of a Laboratory-Confirmed Case of the Coronavirus Disease 2019 (COVID-19)[J].).

34. J. A. Plante et al., Spike mutation D614G alters SARS-CoV-2 fitness. Nature 592, 116–121 (2021).

35. J. A. Backer, D. Klinkenberg, J. Wallinga, Incubation period of 2019 novel coronavirus (2019-nCoV) infections among travellers from Wuhan, China, 20-28 January 2020. Euro Surveill 25, (2020).

36. Q. Li et al., Early Transmission Dynamics in Wuhan, China, of Novel Coronavirus-Infected Pneumonia. N Engl J Med 382, 1199–1207 (2020).

37. B. Rai, A. Shukla, L. K. Dwivedi, Incubation period for COVID-19: a systematic review and meta-analysis. Z Gesundh Wiss, 1–8 (2021).

38. M. Marks et al., Transmission of COVID-19 in 282 clusters in Catalonia, Spain: a cohort study. Lancet Infect Dis, (2021).

39. J. C. Brown, Goldhill, D.H., Zhou, J., Peacock, T.P., Frise, R., Goonawardane, N., Baillon, L., Kugathasan, R., Pinto, A.L., McKay, P.F., Hassard, J., Moshe, M., Singanayagam, A., Burgoyne, T., the ATACCC Investigators, PHE Virology Consortium, Barclay, W.S., Increased transmission of SARS-CoV-2 lineage B.1.1.7 (VOC2020212/01) is not accounted for by a replicative advantage in primary airway cells or antibody escape. BioRxiv, (2021).

40. Y. J. Hou et al., SARS-CoV-2 D614G variant exhibits efficient replication ex vivo and transmission in vivo. Science 370, 1464–1468 (2020).

41. B. Zhou et al., SARS-CoV-2 spike D614G change enhances replication and transmission. Nature 592, 122–127 (2021).

42. Y. Liu et al., The N501Y spike substitution enhances SARS-CoV-2 transmission. bioRxiv, (2021).

43. B. Lubinski, Tang, T., Daniel, S., Jaimes, J.A., Whittaker, G.R., Functional evaluation of proteolytic activation for the SARS-CoV-2 variant B.1.1.7: role of the P681H mutation. bioRxiv 2021.04.06.438731; doi: https://doi.org/10.1101/2021.04.06.438731, (2021).

44. X. Lin et al., ORF8 contributes to cytokine storm during SARS-CoV-2 infection by activating IL-17 pathway. iScience 24, 102293 (2021).

45. A. Stukalov et al., Multilevel proteomics reveals host perturbations by SARS-CoV-2 and SARS-CoV. Nature, (2021).

46. J. F. Pittet et al., TGF-beta is a critical mediator of acute lung injury. Journal of Clinical Investigation 107, 1537–1544 (2001).

47. Y. Aschner, G. P. Downey, Transforming Growth Factor-beta: Master Regulator of the Respiratory System in Health and Disease. Am J Respir Cell Mol Biol 54, 647–655 (2016).

48. P. Domingo-Calap, E. Segredo-Otero, M. Duran-Moreno, R. Sanjuan, Social evolution of innate immunity evasion in a virus. Nat Microbiol 4, 1006–1013 (2019).

49. R. Challen et al., Risk of mortality in patients infected with SARS-CoV-2 variant of concern 202012/1: matched cohort study. BMJ 372, n579 (2021).

50. T. Sato et al., Long-term expansion of epithelial organoids from human colon, adenoma, adenocarcinoma, and Barrett’s epithelium. Gastroenterology 141, 1762–1772 (2011).

51. J. Beumer et al., Enteroendocrine cells switch hormone expression along the crypt-to-villus BMP signalling gradient. Nat Cell Biol 20, 909–916 (2018).

52. V. M. Corman et al., Detection of 2019 novel coronavirus (2019-nCoV) by real-time RT-PCR. Euro Surveill 25, (2020).

53. B. B. Oude Munnink et al., Rapid SARS-CoV-2 whole-genome sequencing and analysis for informed public health decision-making in the Netherlands. Nat Med 26, 1405–1410 (2020).

54. M. Martin, Cutadapt removes adapter sequences from high-throughput sequencing reads. EMBnet.journal 17, (2011).

55. B. Langmead, S. L. Salzberg, Fast gapped-read alignment with Bowtie 2. Nat Methods 9, 357–359 (2012).

56. G. Jun, M. K. Wing, G. R. Abecasis, H. M. Kang, An efficient and scalable analysis framework for variant extraction and refinement from population-scale DNA sequence data. Genome Research 25, 918–925 (2015).

57. D. C. Koboldt et al., VarScan 2: somatic mutation and copy number alteration discovery in cancer by exome sequencing. Genome Res 22, 568–576 (2012).

58. H. Li et al., The Sequence Alignment/Map format and SAMtools. Bioinformatics 25, 2078–2079 (2009).

59. A. Tareen, J. B. Kinney, Logomaker: beautiful sequence logos in Python. Bioinformatics 36, 2272–2274 (2020).

60. T. Hashimshony et al., CEL-Seq2: sensitive highly-multiplexed single-cell RNA-Seq. Genome Biol 17, 77 (2016).

61. S. Simmini et al., Transformation of intestinal stem cells into gastric stem cells on loss of transcription factor Cdx2. Nat Commun 5, 5728 (2014).

62. H. Li, R. Durbin, Fast and accurate long-read alignment with Burrows-Wheeler transform. Bioinformatics 26, 589–595 (2010).

63. M. I. Love, W. Huber, S. Anders, Moderated estimation of fold change and dispersion for RNA-seq data with DESeq2. Genome Biol 15, 550 (2014).

64. F. G. Faas et al., Virtual nanoscopy: generation of ultra-large high resolution electron microscopy maps. J Cell Biol 198, 457–469 (2012).

